# An expanded analysis framework for multivariate GWAS connects inflammatory biomarkers to functional variants and disease

**DOI:** 10.1101/867267

**Authors:** Sanni E. Ruotsalainen, Juulia J. Partanen, Anna Cichonska, Jake Lin, Christian Benner, Ida Surakka, FinnGen, Mary Pat Reeve, Priit Palta, Marko Salmi, Sirpa Jalkanen, Ari Ahola-Olli, Aarno Palotie, Veikko Salomaa, Mark J. Daly, Matti Pirinen, Samuli Ripatti, Jukka Koskela

**Author notes:** authors contributed equally (Sanni E. Ruotsalainen, Juulia J. Partanen).

## Abstract

Multivariate methods are known to increase the statistical power of association detection, but they have lacked essential follow-up analysis tools necessary for understanding the biology underlying these associations. We developed a novel computational workflow for multivariate GWAS follow-up analyses, including fine-mapping and identification of the subset of traits driving associations (driver traits). Many follow-up tools require univariate regression coefficients which are lacking from multivariate results. Our method overcomes this problem by using Canonical Correlation Analysis to turn each multivariate association into its optimal univariate Linear Combination Phenotype (LCP). This enables an LCP-GWAS, which in turn generates the statistics required for follow-up analyses. We implemented our method on 12 highly correlated inflammatory biomarkers in a Finnish population-based study. Altogether, we identified 11 associations, four of which (*F5, ABO, C1orf140 and PDGFRB*) were not detected by biomarker-specific analyses. Fine-mapping identified 19 signals within the 11 loci and driver trait analysis determined the traits contributing to the associations. A phenome-wide association study on the 19 putative causal variants from the signals in 176,899 individuals from the FinnGen study revealed 53 disease associations (p < 1×10^-4^). Several reported pQTLs in the 11 loci provided orthogonal evidence for the biologically relevant functions of the putative causal variants. Our novel multivariate analysis workflow provides a powerful addition to standard univariate GWAS analyses by enabling multivariate GWAS follow-up and thus promoting the advancement of powerful multivariate methods in genomics.

## INTRODUCTION

Genome-wide association studies (GWAS) of biomarkers have been highly successful in identifying novel biological pathways and their impact on health and disease. Biomarkers increase statistical power in GWAS, compared to disease diagnoses, due to their quantitative nature and lack of errors due to subjectivity, such as misclassification. Thus, biomarker GWAS have identified thousands of biomarker-associated loci and elucidated the mechanisms underlying numerous disease associations^1–3^. A recent study on 38 biomarkers in the UK Biobank (UKBB) identified over 1,800 independent genetic associations with causal roles in several diseases^4^. Proteomics and metabolomics integrated with genomics has also revealed causal molecular pathways connecting the genome to multiple diseases, e.g. autoimmune disorders and cardiovascular disease^5–8^. Although biomarkers are more closely related to pathophysiology, a single biomarker is usually an inaccurate estimator of complex disease due to phenotypic heterogeneity and individual variation. Therefore, combinations of biomarkers provide a more robust predictive molecular signature. Studies examining combinations of biomarkers are increasingly feasible given the availability of biobank resources around the globe with deep phenotyping, i.e. precise and comprehensive data on phenotypic variation including quantitative measures such as biomarkers^9, 10^.

Multivariate GWAS of correlated traits increases statistical power compared to univariate analysis, especially in the case of complex biological processes and correlated traits^8, 11, 12^. This leads to identifying multivariate associations that are otherwise missed by univariate analysis^8, 13^. Efficient software programs are available for performing multivariate GWAS such as metaCCA^14^, yet multivariate analyses currently have shortcomings in interpreting the arising signals. Follow-up tools for fine-mapping causal variants within the associated loci are lacking and the subset of tested traits that drive the association signals have not been identified. These shortcomings are largely due to the lack of a multivariate counterpart to the univariate regression coefficients (beta estimates). Lack of these necessary follow-up tools has hindered the utilization of multivariate methods.

In this study, we developed a novel computational workflow for multivariate GWAS discovery and follow-up analyses including fine-mapping and identification of driver traits (Figure 1). Our workflow includes 1) a customized version of the metaCCA software that overcomes the problem of missing beta estimates by turning each multivariate association into its optimal univariate Linear Combination Phenotype (LCP), enabling an LCP-GWAS, 2) fine-mapping, i.e. identifying putative causal variants underlying each association using summary statistics from the LCP-GWAS and a multivariate extension to FINEMAP^15^, and 3) determining the traits driving each multivariate association using a newly developed tool, MetaPhat^16^ that efficiently decomposes the multivariate associations into a smaller set of underlying driver traits. Taken together, we present to our knowledge the first comprehensive framework to map multivariate associations into individual causal variants and a subset of driver traits. We demonstrate the potential of our workflow in a Finnish population-based cohort with 12 inflammatory biomarkers implicated in the pathogenesis of autoimmune disorders and cancer^17–19^. This set of highly-correlated biomarkers is particularly advantageous for multivariate analysis as high correlation between traits increases the boost in statistical power achieved by multivariate methods. Using multivariate analysis, we identify additional hits compared to univariate analysis, totaling 11 independent associations. We follow them up in a phenome-wide association study (PheWAS) in the FinnGen study (n = 176,899) across 2,367 disease endpoints and in the UKBB (n = 408,910) ^10^. We discover multiple disease associations, as well as identify orthogonal evidence for the biological impact of the causal variants through several expression quantitative trait loci (eQTLs) and protein quantitative trait loci (pQTLs) within the multivariate loci.

**Figure 1.**
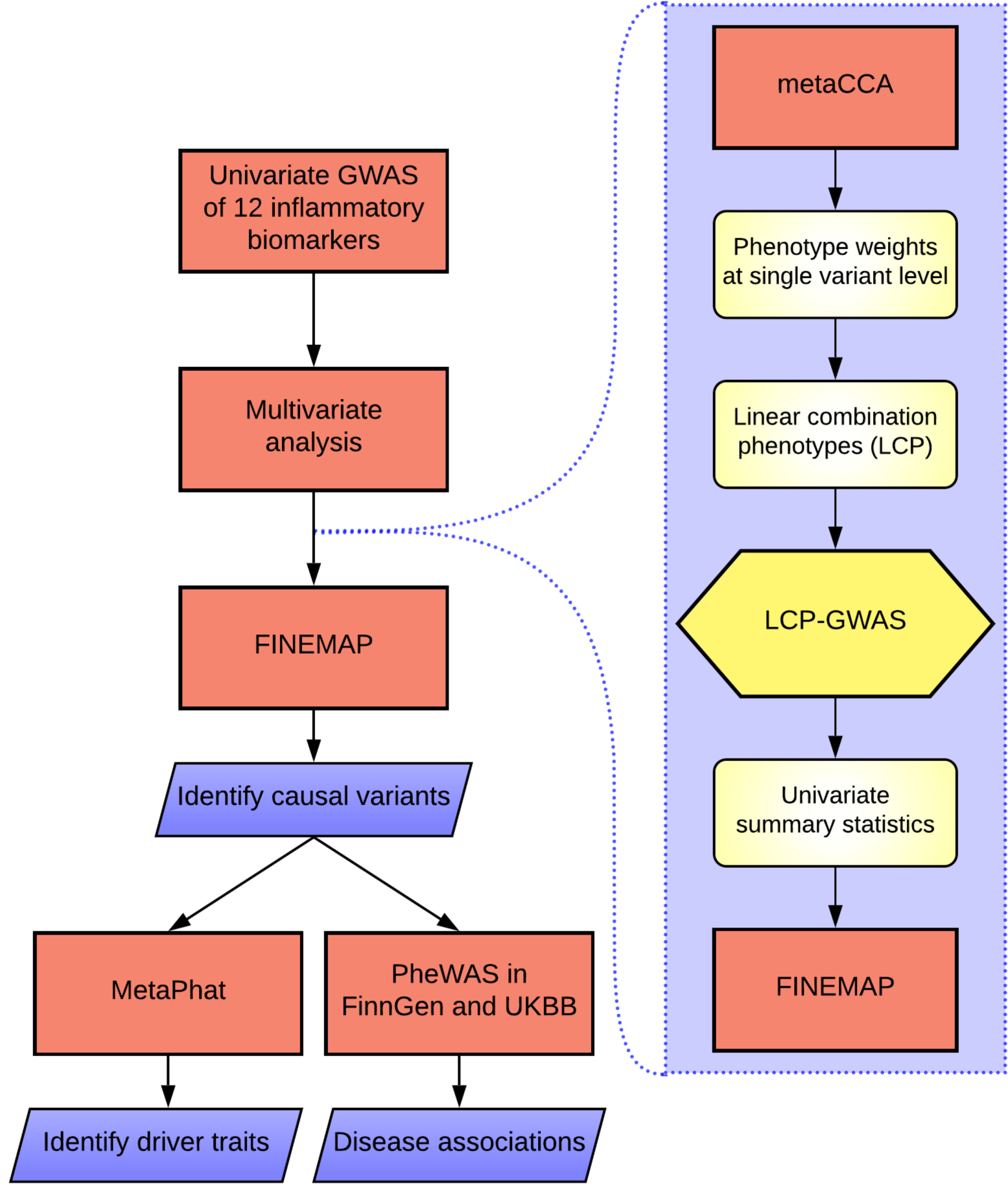
Study workflow. The novel LCP-GWAS method that enables follow-up analyses such as fine-mapping for multivariate GWAS is illustrated in the violet panel on the right.

## MATERIALS AND METHODS

### Study cohort and data

We studied 12 highly correlated inflammatory biomarkers in the population-based national FINRISK Study collected in 1997 (n = 6,890) ^20^ (Table 1, Supplementary Figure 1). The FINRISK Study is a large Finnish population survey of risk factors for chronic, non-communicable diseases, and it has been collected by independent random population sampling every five years beginning in 1972 with multiple recruiting waves. The 12 inflammatory biomarkers included five interleukins (IL-4, IL-6, IL-10, IL-12p70, IL-17), three growth factors (FGF2, PDGF-BB, VEGF-A), one colony-stimulating factor (G-CSF), one interferon (IFN-γ), one chemokine (SDF-1ɑ), and one tumor necrosis factor (TNF-β) (Table 1, Supplementary Figure 1. Hierarchical clustering identified the cluster of 12 inflammatory biomarkers out of 66 quantitative traits of cardiometabolic or immunologic relevance (Supplementary Figure 2 Supplementary Table 1 and Supplementary Methods). The 66 quantitative traits were measured as previously described^11, 20, 21^.

**Table 1.**
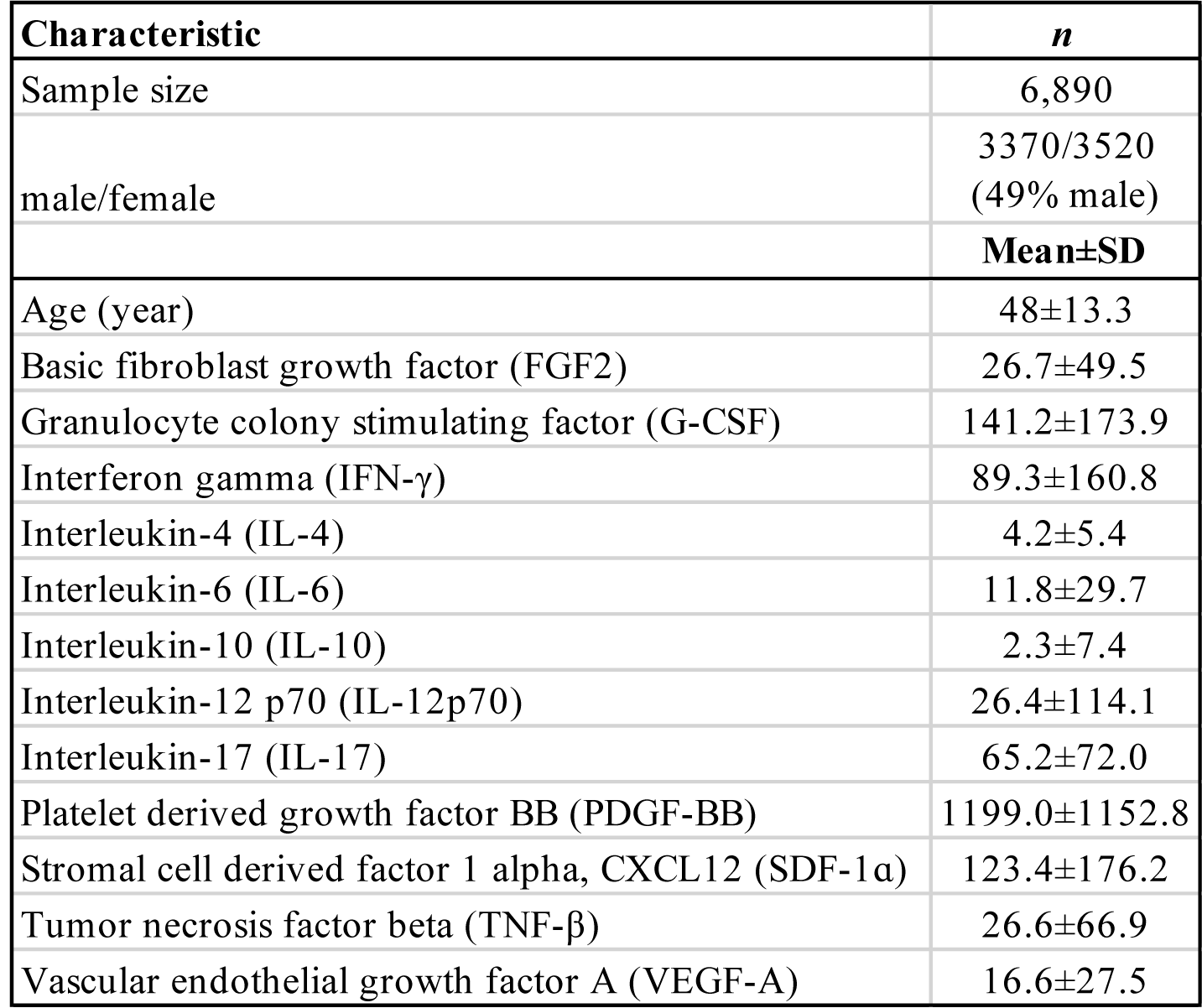
Characterization of the 12 inflammatory biomarker measurements. n = sample size, SD = standard deviation. The cytokine concentrations are pg/ml.

### Genotyping, imputation and quality control

Samples were genotyped using multiple different genotyping chips, for which the quality control (QC), phasing and imputation were done in multiple chip-wise batches. Imputation of the genotypes was done utilizing a Finnish population-specific reference panel of 3,775 high-coverage whole-genome sequences. Genotype imputation was followed by post-imputation sample QC (Supplementary Methods) and variant QC (imputation INFO > 0.8, minor allele frequency > 0.002 and Hardy-Weinberg equilibrium p-value > 1×10^-6^). A total of 26,717 samples and 11,329,225 variants passed this rigorous quality control.

### Univariate and multivariate GWAS

Univariate genome-wide association analyses for the biomarkers were performed using a linear mixed model implemented in Hail^22^, adjusting for age, sex, genotyping chip, first ten principal components of genetic structure and the genetic relationship matrix (GRM) (Supplementary Methods). The GRM was estimated using 73K independent high-quality genotyped variants (Supplementary Methods). We performed multivariate GWAS on the biomarkers using metaCCA^14^, software that performs multivariate analysis by implementing Canonical Correlation Analysis (CCA) for a set of univariate GWAS summary statistics.

The objective of CCA is to find the linear combination of the *p* predictor variables (*X_1_, X_2_, …, X_p_*) that is maximally correlated with a linear combination of the *q* response variables (*Y_1_, Y_2_, …, Y_q_*). If we denote the respective linear combinations by

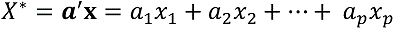

and

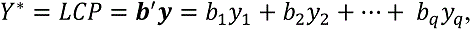

then finding the linear combination of the predictor variables that are maximally correlated with the linear combination of the response variables corresponds to finding vectors **a** and **b** that maximize

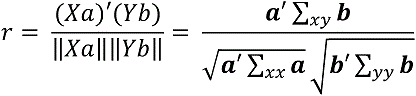

where Σ_xx_, Σ_yy_ and Σ_xy_ represent the variance-covariance matrices of the predictor variables, response variables and both of them together, respectively. The maximized correlation *r* is the *canonical correlation* between **X** and **Y**. Multivariate GWAS is a special case of CCA with multiple response variables *Y*, but only one explanatory variable *X*, the genotypes at the variant tested.

### Novel multivariate LCP-GWAS method

To enable follow-up analyses of multivariate GWAS results, such as fine-mapping, we developed a novel method to produce linear combination phenotypes (LCP) at the single variant level by extending the functionality of metaCCA. The updated metaCCA is available online at: https://github.com/acichonska/metaCCA.

LCPs were constructed as the weighted sum of the trait residuals, where the weights (***b*** = [*b_1_, b_2_ …, b_q_*]) were chosen to maximize the correlation between the resulting linear combination of traits and the genotypes at the variant. We determined association regions by adding 1Mb to each variant reaching genome-wide significance (GWS; p-value < 5×10^-8^) in the multivariate analysis and joining overlapping regions. We constructed LCPs for the lead variant, i.e. the variant with the smallest p-value, in each of these regions, as a univariate representation of the multivariate association in that region. Next, we performed chromosome-wide LCP-GWAS for the constructed LCPs in a similar manner as for each of the biomarkers.

### Fine-mapping multivariate associations

We used FINEMAP^15, 23^ on the LCP-GWAS summary statistics to identify causal variants underlying the multivariate associations. FINEMAP analyses were restricted to a ±1Mb region around the GWS variants from the LCP-GWAS.

We assessed variants in the top 95% credible sets, i.e. the sets of variants encompassing at least 95% of the probability of being causal (causal probability) within each causal signal conditional on other causal signals in the genomic region. Within these sets we excluded those sets that did not clearly represent one signal, determined by low minimum linkage disequilibrium (LD, r^2^ < 0.1).

To validate the multivariate fine-mapping results, we also performed conventional stepwise conditional analysis for all fine-mapping regions using LCPs. We iteratively conditioned on the lead variant in the region until the smallest p-value in the region exceeded 5×10^-8^.

### Identifying driver traits

We determined the traits driving the multivariate associations for the putative causal variants suggested by fine-mapping using the MetaPhat software developed in-house^16^. MetaPhat determines the set of driver traits for each multivariate association by performing multivariate testing using metaCCA iteratively on subsets of the traits, excluding one trait at a time until a single trait remains. At each iteration, the trait to be excluded is the one whose exclusion leads to the highest p-value for the remaining subset of traits. The driver traits are determined as a set of traits that have been removed when the multivariate p-value becomes non-significant (p > 5×10^-8^). The interpretation is that the driver traits make the multivariate association significant.

### Phenome-wide association testing in FinnGen and UKBB

We performed a PheWAS in the FinnGen study for variants suggested to be causal by multivariate fine-mapping and for multivariate GWS functional variants (Table 2 and Supplementary Table 2. FinnGen (https://www.finngen.fi/en) is a large biobank study that aims to genotype 500,000 Finns and combine this data with longitudinal registry data, including national hospital discharge, death, and medication reimbursement registries, using unique national personal identification numbers. FinnGen includes prospective epidemiological and disease-based cohorts as well as hospital biobank samples. A total of 176,899 samples from FinnGen Data Freeze 4 with 2,444 disease endpoints were analyzed using Scalable and Accurate Implementation of Generalized mixed model (SAIGE), which uses saddlepoint approximation (SPA) to calibrate unbalanced case-control ratios^24^. Additional details and information on the genotyping and imputation are provided in the Supplementary Material and contributors of FinnGen are listed in the Acknowledgements.

**Table 2.**
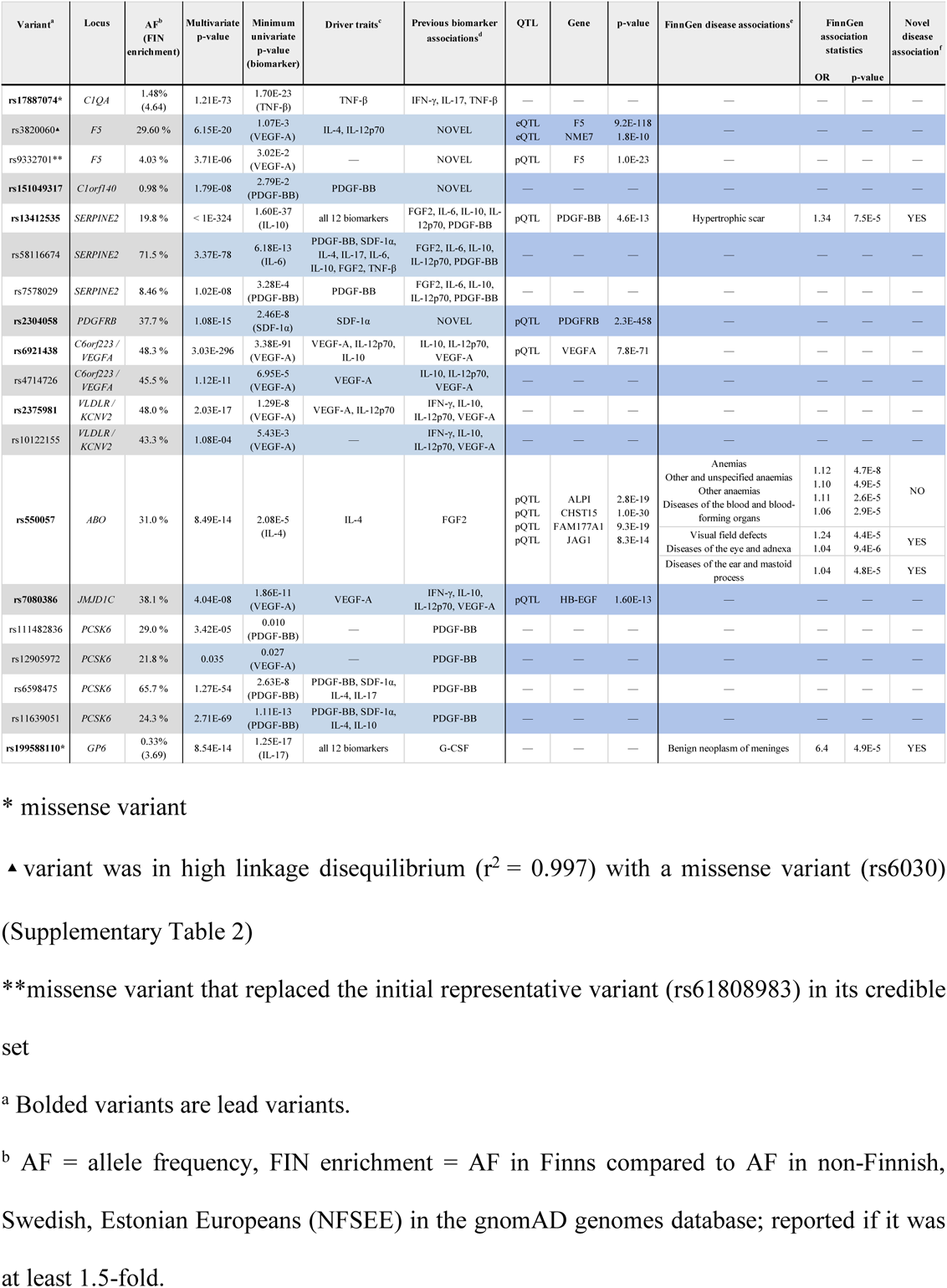
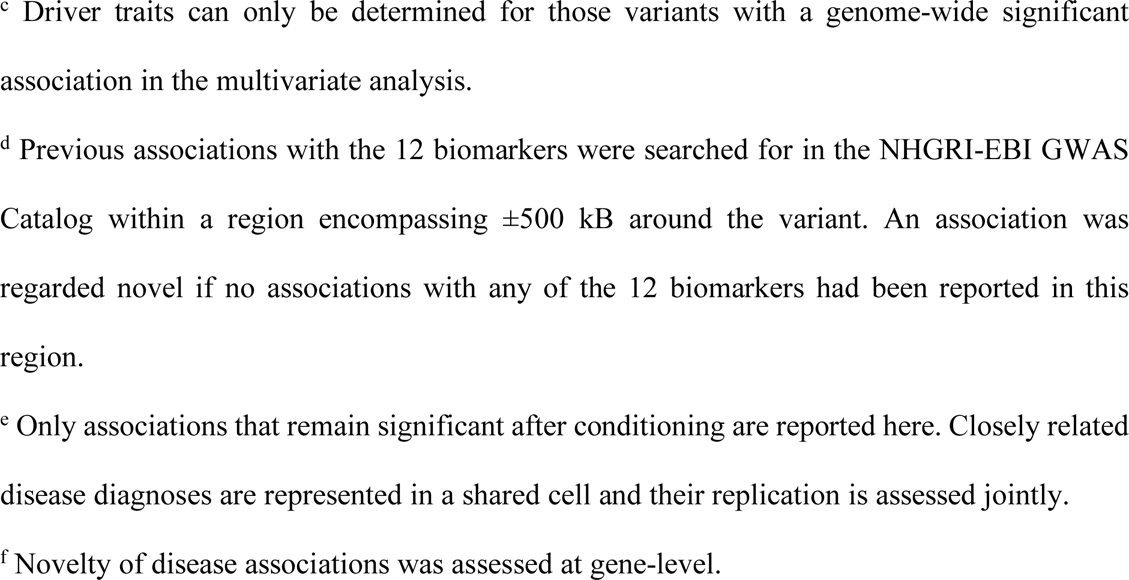
Results of the 19 putative causal variants, i.e. the representative variants of the 19 credible sets.

FinnGen disease associations with p-values < 1×10^-4^ were considered significant. We tested the p-value threshold by sampling 1,000 allele frequency-matched sets of *n* variants, where *n* represents the number of variants of interest, from 8.2 million non-coding variants and determining a null distribution of the number of FinnGen associations passing the p-value threshold. We confirmed the validity of the p-value threshold by comparing the observed number of FinnGen associations passing the p-value threshold to the null distribution (Supplementary Figure 3). We excluded disease endpoints within the ICD-10 (International Statistical Classification of Diseases and Related Health Problems 10th Revision) chapters XXI and XXII from PheWAS analyses, resulting in 2,367 disease endpoints analyzed. To assess the relevance of the putative causal variants and the functional variants for their disease associations in FinnGen, the disease associations were conditioned on the variant with the strongest FinnGen disease association within the locus (±0.5MB of the putative causal variant or functional variant). Finally, we assessed replication of the disease associations in the UKBB, where associations with p-values < 0.05 were considered replicated given that the direction of effects were coherent. Phecodes from the UKBB were mapped to ICD-10 diagnosis codes using the PheCode map 1.2^25^. The NHGRI-EBI GWAS Catalog^26^ was used for assessing the novelty of the observed genetic associations.

We also explored whether the fine-mapped putative causal variants or variants in LD with them (r^2^ > 0.6) had previously been reported as eQTLs or pQTLs. For eQTLs, we looked at overlap with LD-pruned associations derived from the Genotype-Tissue Expression (GTEx) Portal and pQTLs were included from studies by Suhre^5^, Sun^6^, Emilsson^27^ and Sasayama^28^; regional overlap and architecture were visualized in Target Gene Notebook^29^.

## RESULTS

### Comparison of multivariate and univariate GWAS of 12 inflammatory biomarkers

We first tested for genome-wide associations of 12 highly correlated inflammatory biomarkers (Table 1, Supplementary Figure 1) measured in 6,890 FINRISK study participants using both multivariate and univariate methods. Pearson correlations between the biomarkers ranged from 0.64 to 0.93, with a mean of 0.80. Out of the 11,329,225 variants tested, 190 were significantly associated using both univariate and multivariate analyses, 999 (in 11 loci) only by the multivariate analysis and two only by the univariate analysis using a Bonferroni-corrected p-value threshold of 5×10^-8^/12 (Figure 2). The two variants that were significant only in the univariate analysis were both located in a locus (*JMJD1C*) that was found to be significant also by the multivariate analysis. A total of 1,189 variants reached the significance threshold in the multivariate analysis compared to only 192 in the univariate analysis, reflecting a considerable increase in statistical power achieved by the multivariate analysis while preserving the Type I error rate (Supplementary Figure 4). The corresponding genomic inflation factor λ was 1.036, with no evidence of concerning genomic inflation due to multivariate analysis using Canonical Correlation Analysis.

**Figure 2.**
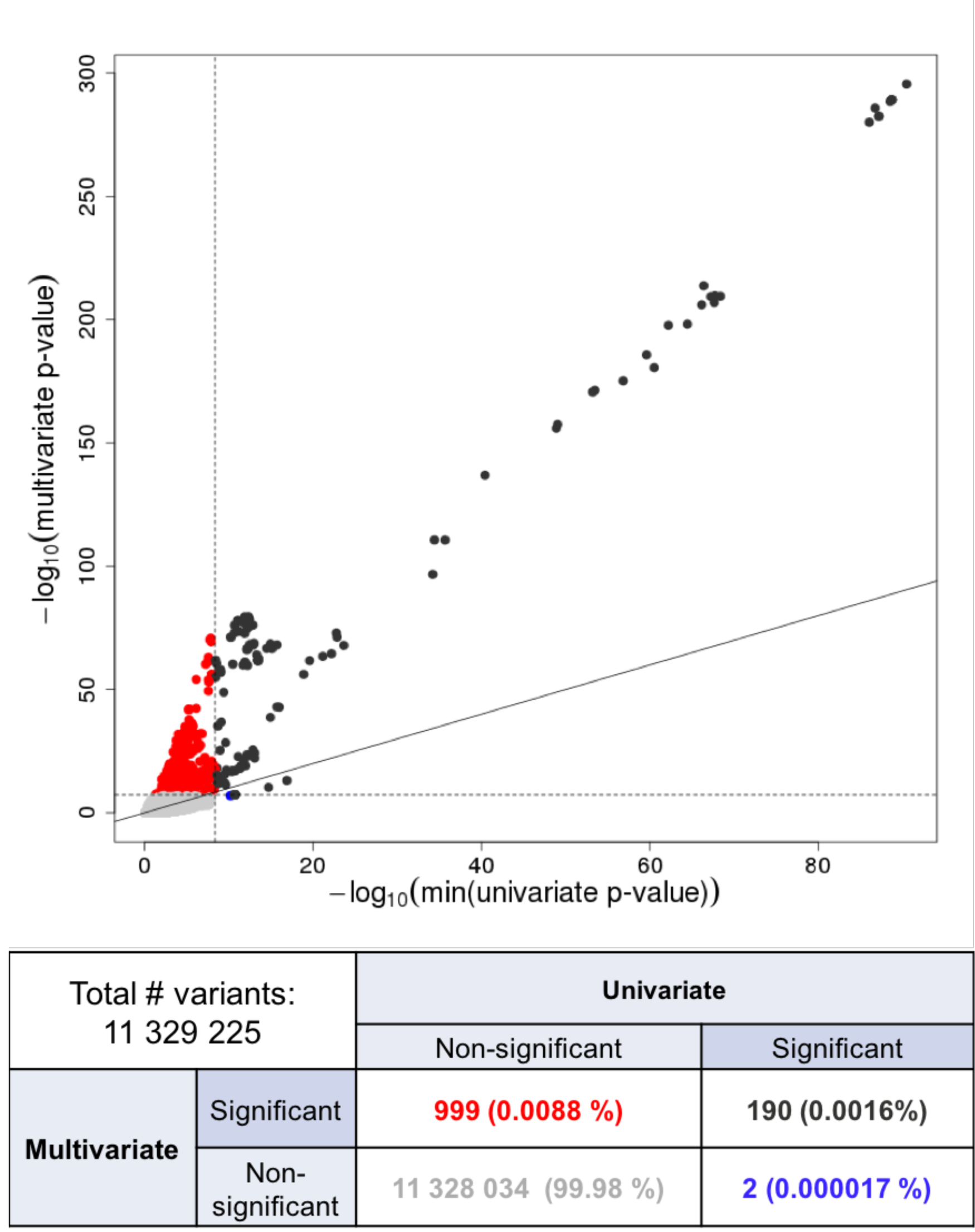
Power comparison between multivariate and univariate methods. Red and blue dots represent genetic variants reaching genome-wide significance only by the multivariate or univariate method, respectively. Black dots reach the genome-wide significance threshold by both methods and grey dots do not by either method. Respective numbers are reported in the accompanying table.

Within the 1,189 genome-wide significant variants in the multivariate analysis, we identified 11 independently associated loci (Figure 3 and Supplementary Figure 5), four of which (*F5, C1orf140, PDGFRB* and *ABO*) were not detected by univariate analyses corrected for multiple testing (Figure 3). Eight of the 11 loci had previously been associated with at least one of the 12 biomarkers in the NHGRI-EBI GWAS catalog while three loci (*F5, C1orf140* and *PDGFRB*) were novel.

**Figure 3.**
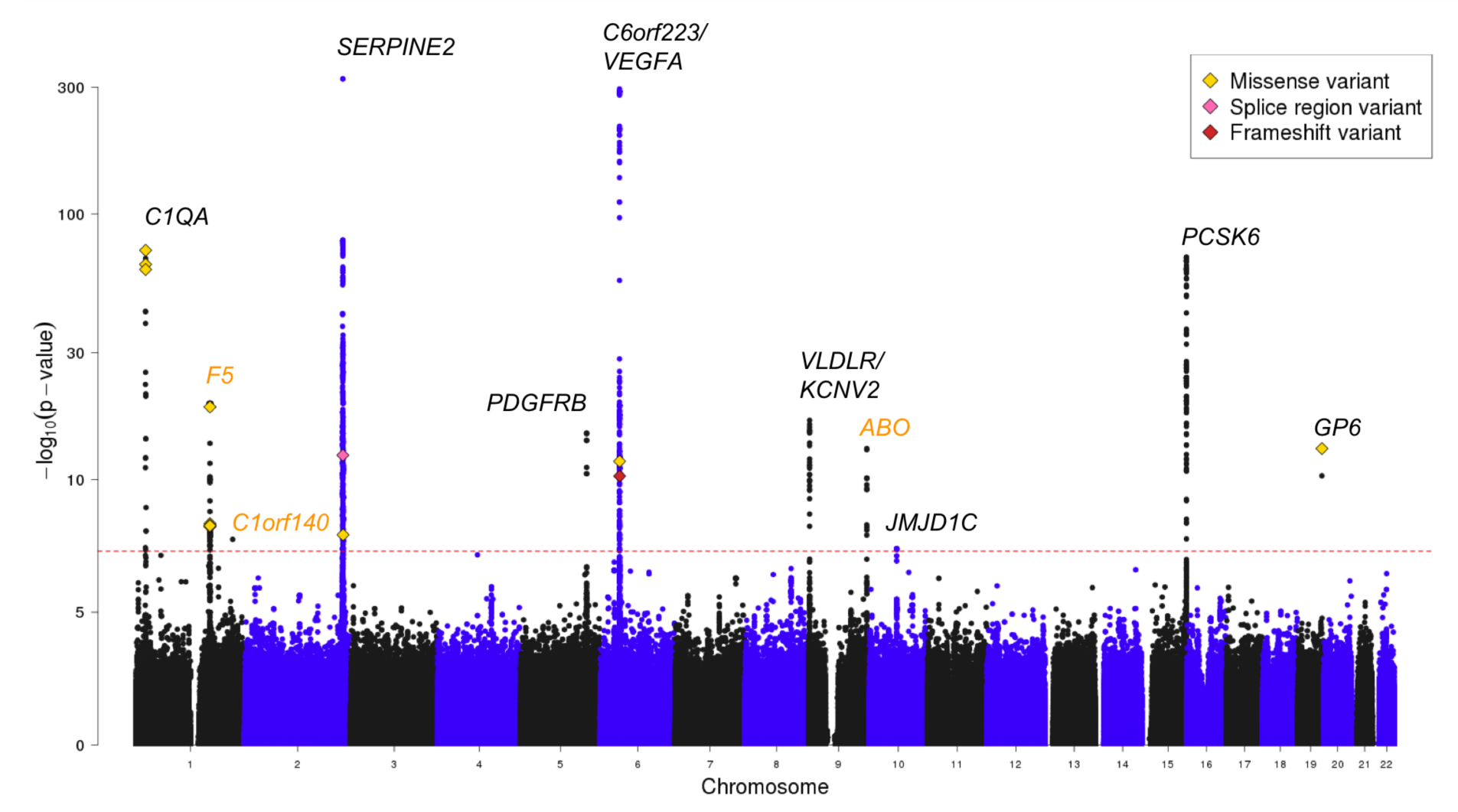
Manhattan plot of the multivariate GWAS results on 12 inflammatory biomarkers. Gene names colored in orange represent associations only detected by the multivariate method while black are detected by both multivariate and univariate methods. 13 genome-wide significant functional variants are denoted with diamonds.

At several loci, multivariate analysis revealed more plausible candidates for causal variants than the univariate analyses. For example, in the *C1QA* locus, an association with only one of the 12 biomarkers, TNF-β, was noted in the univariate results. The lead variant in the TNF-β univariate GWAS was rs78655189 (p = 2.2×10^-24^), an intronic variant in the *EPHB2* gene. In contrast, the lead variant for the same locus in the multivariate analysis was rs17887074 (p = 1.2×10^-73^), a Finnish-enriched missense variant located in the *C1QA* gene. The *C1QA* gene has been previously associated with immunologic diseases, such as immunodeficiency and systemic lupus erythematosus^30^. Our multivariate analysis may point towards a plausible mechanism underlying these associations via TNF-β levels. Further, in the *VLDLR* locus, univariate fine-mapping of VEGF, the only associated biomarker, suggested that the lead variant rs2375981 from the multivariate analysis was more likely causal than the lead variant rs10967570 from the VEGF univariate analysis (posterior probabilities 1.0 and 0.025, respectively).

### Fine-mapping multivariate GWAS results

To identify the causal variants of the multivariate associations, we studied the likelihood of multiple variants contributing to the association signal in the 11 associated loci using FINEMAP^23^. Our novel multivariate LCP-GWAS method based on linear combinations calculated for each locus using multivariate metaCCA results enabled fine-mapping of the multivariate results. The number of credible sets varied from one to four for the multivariate associated loci (Supplementary Table 3), resulting in a total of 19 independent sets of variants considered putatively causal. All 183 variants within the 19 credible sets are available in Supplementary Table 3 and posterior probabilities for different numbers of causal signals for each locus are available in Supplementary Table 4.

Among each of the 19 sets, the variant with the highest the causal probability (initial representative variant) was chosen to represent the set, unless the set contained a functional variant (missense, splice-region or frameshift) in high LD (r^2^ > 0.95) with the initial representative variant, in which case the functional variant was chosen (Table 2 and Supplementary Figure 6). This was the case for one credible set in the *F5* locus where the missense variant rs9332701 (causal probability 46.1%) replaced the initial representative non-coding variant rs61808983 (causal probability 53.3%) as they were in high LD (r^2^ = 0.996). We also assessed whether the causal probabilities changed in the *F5* credible set if the LCP was generated for the missense variant rs9332701 rather than the lead variant rs61808983. This had no notable effects on the causal probabilities (46.1% vs. 48.5%, 53.3% vs. 51.5% for rs9332701 and rs61808983, respectively).

The 19 representative variants, hereon referred to as the putative causal variants, included all except one (rs11637184 in *PCSK6* locus) of the 11 lead variants from multivariate GWAS. In the *PCSK6* locus one of the four putative causal variants (rs111482836) was associated with disease in FinnGen, whereas the lead variant was not, highlighting the importance of fine-mapping multivariate GWAS results.

Fine-mapping suggested at least as many causal signals as there were conditional rounds in stepwise conditional analysis (n = 16), thus verifying the results from FINEMAP. Further, 13 of the 19 (68,4%) putative causal variants were also conditioned on in the conditional analysis (Supplementary Table 5). The main benefit of fine-mapping is the probabilistic quantification of causality for each variant in the region, which is crucial information when there are several plausible candidates for causal variants. Such metrics are not available from stepwise conditional analysis.

### Functional coding variants

GWAS hits are generally non-coding, although concentrated in regulatory regions^31^, and enrichment of functional coding variants has been seen mainly only after fine-mapping e.g. in inflammatory bowel disease^32^. We, however, observed enrichment of functional coding variants in the multivariate GWAS hits already prior to fine-mapping. Two of the 19 putative causal variants were missense variants (rs17887074 and rs199588110, in the *C1QA* and *GP6* loci respectively). These two variants (2/19, 10.5%) were enriched (>1.5-fold) in Finns compared to non-Finnish, Swedish, Estonian

Europeans (NFSEE) in the gnomAD genome reference database^33^. Considering all genome-wide significant variants in the multivariate GWAS, we found 13 functional variants (missense, splice-region and frameshift variants) with at least one functional variant in five of the 11 multivariate loci (*C1QA, F5, C1orf140, SERPINE2,* and *GP6*; Supplementary Table 2). Out of the 13 functional variants, 11 were missense variants, one was a splice region variant and one a frameshift variant. A total of six (46.2%) of the 13 variants were enriched in the Finnish population, highlighting the potential of utilizing isolated populations in GWAS.

We studied whether the multivariate genome-wide significant variants and variants identified by fine-mapping were enriched for functional variants including missense, splice-region and frameshift variants compared to the 11.3M variants analyzed. P-values for enrichment were calculated using the *X*^2^-test for the number of functional or missense variants within the variants assessed against the number of the corresponding subset of variants within all variants tested. The multivariate genome-wide significant variants were enriched for missense variants and functional variants including missense, splice-region and frameshift variants (2.2-fold, p = 0.015, and 1.9-fold, p = 8.8×10^-4^, respectively). The 19 putative causal variants were further enriched for both missense variants and the broader set of functional variants (37-fold, p = 1.3×10^-17^, and 28-fold, p = 1.4×10^-17^, respectively) as were the 183 variants in the credible sets (3.9-fold, p = 0.050, and 2.9-fold, p = 0.050, respectively).

### Identifying driver traits

Next, we studied which traits were driving the multivariate associations in each of the 11 loci using metaPhat^16^. The number of driver traits for each of the 11 loci varied between one and all 12. The driver traits were very much in line with the univariate results; the most significantly associated biomarkers in the univariate GWAS were typically included among the driver traits (Table 2). In loci with multiple putative causal variants, driver traits for the variants were generally subsets of the lead variant’s driver traits, and a stronger multivariate association increased the number of driver traits. However, this relationship between multivariate p-value and the number of driver traits did not hold across loci. Further, driver traits typically included all or some of the biomarkers that had previously been associated with the locus (Table 2).

### Disease implications of the multivariate loci

Finally, we tested how the 19 putative causal variants as well as the 13 genome-wide significant functional variants in the 11 loci associated with disease risk among 2,367 disease endpoints defined in FinnGen. Altogether, 53 disease associations were observed with seven putative causal variants. Two of these variants did not lead the multivariate associations at the 11 loci and thus would have gone unnoticed without fine-mapping. Five genome-wide significant functional variants not overlapping with the putative causal variants had an additional 35 disease associations.

To assess the relevance of the putative causal variants and the functional variants for their disease associations in FinnGen, the disease associations were conditioned on the variant with the strongest FinnGen disease association within the locus. In 13 of the 53 FinnGen disease associations with the putative causal variants, the putative causal variant or a variant in near perfect LD (r^2^ > 0.95) led the association signal or remained significant after conditioning. Correspondingly, for the functional variants not overlapping with the putative causal variants 18 of the 35 disease associations were either led by the functional variant or a variant in near perfect LD or remained significant after conditioning. We also tested the disease associations in the UKBB, where associations with p-values < 0.05 were considered replicated given that the direction of effects were coherent (Supplementary Table 6).

In addition to disease associations, we explored whether the putative causal variants or variants in LD with them (r^2^ > 0.6) had previously been reported as eQTLs or pQTLs. Several reported eQTLs and pQTLs^5^ in the 11 loci provided orthogonal evidence for the biologically relevant functions of the putative causal variants (Supplementary Table 7).

Here we further discuss results for the four multivariate loci with disease associations (p < 1×10^-4^) in FinnGen that remained significant after conditioning. The variants identified by multivariate testing for which the associations became insignificant after conditioning, were regarded unnecessary for the observed disease association. Full disease association results for the 11 loci are shown in Supplementary Table 8.

### *GP6* gene locus

#### Multivariate association and FinnGen disease associations

The Finnish enriched rare missense variant rs199588110 (AF = 0.33%, 3.7-fold enrichment), predicted deleterious by SIFT^34^ and probably damaging by Polyphen^35^, was suggested causal in the *GP6* locus. In FinnGen it led the association with benign neoplasms of meninges (OR = 6.4, p = 4.9×10^-5^). The association was not replicated in the UKBB, although this may be due to impaired power as the AF of the Finnish enriched variant in the UKBB (0.036%) was roughly a tenth of its AF in FinnGen, and an inadequate match of the discovery and replication phenotypes as UKBB phenotype definitions included all benign neoplasms of the brain and spinal cord and were not restricted to neoplasms of the meninges.

#### Driver traits

All 12 biomarkers were considered driver traits of the multivariate association. Cytokines, including many of the 12 biomarkers studied (e.g. IL-6, IL-4, PDGF-BB and VEGF-A), have been implicated in the autocrine regulation of meningioma cell proliferation and motility^36–39^. Further, higher expression levels of both PDGF-BB and VEGF occur in atypical and malignant meningiomas than in benign meningiomas^39, 40^ and microvascular density regulated by VEGF has been linked with time to recurrence^41^. Several phase II clinical trials have tested therapies targeting VEGF and PDGF-BB signaling pathways as treatments for recurrent or progressive meningiomas^37^ with promising results for two multifunctional tyrosine kinase inhibitors, sunitinib and PTK787/ZK 222584 that inhibit both VEGF and PDGF receptors^37, 42^.

### *SERPINE2* gene locus

#### Multivariate association and FinnGen disease associations

The *SERPINE2* locus was the locus with the most significant association in the multivariate analysis (p < 1×10^-324^). Three variants (rs13412535, rs58116674 and rs7578029) were suggested causal (putative causal variants). One of them, the intronic lead variant rs13412535 from the multivariate analysis, increased the risk of hypertrophic scars (OR = 1.3, p = 7.5×10^-5^) and was in very high LD with the variant that led the disease association in FinnGen (rs68066031, r^2^ = 0.99). The association was not replicated in the UKBB and had not been previously reported at gene-level. Nonetheless, the variant in question had an association with another hypertrophic skin disorder, acquired keratoderma (OR = 1.5, P = 0.02) in the UKBB.

#### Previous knowledge of gene function and driver traits

The *SERPINE2* gene encodes protease nexin-1, a protein in the serpin family of proteins that inhibits serine proteases, especially thrombin, and has therefore been implicated in coagulation and tissue remodeling^43^. The gene has been associated with chronic obstructive pulmonary disease and emphysema^44^. As previously reported, *SERPINE2* has been shown to inhibit extracellular matrix degradation^45^ and overexpression of *SERPINE2* has been shown to contribute to pathological cardiac fibrosis in mice^46^. Additionally, serine protease inhibitor genes including *SERPINE2* have been noted to be heavily induced during wound healing^47^. According to GTEx the *SERPINE2* gene is most highly expressed in fibroblasts. Further, inflammation plays an important role in hypertrophic scar formation and cytokines including PDGF and VEGF are dysregulated in hypertrophic scars^48^. The lead variant had genome-wide significant associations with 11 of the 12 biomarkers and all 12 were regarded as driver traits of the association.

#### eQTLs and pQTLs

The lead variant (rs13412535) is a pQTL impacting one of the driver traits, PDGF-BB levels, and an intronic variant rs68066031 in high LD (r^2^ = 0.99) with the lead variant is a pQTL for SERPINE2^6, 27^. PDGF is considered essential in wound repair^49^ and growth factors including PDGF are considered key players in the pathogenesis of hypertrophic scars^50^. PDGF enhances pathologic fibrosis in several tissues such as skin, lung, liver and kidney by means of mitogenic and chemoattractant actions on the principal collagen-producing cell type, myofibroblasts, as well as stimulation of collagen production^51^.

### *ABO* gene locus

#### Multivariate association and FinnGen disease associations

An association with the *ABO* locus was only detected by multivariate analysis (minimum univariate p = 2.1×10^-5^ for the lead variant from multivariate analysis). One variant, the intronic lead variant rs550057 (aka rs879055593) from multivariate analysis (p = 8.5×10^-14^) was suggested causal and was associated with 45 endpoints in FinnGen, such as endometriosis, heart failure and statin usage. Most of these associations resulted from LD to other stronger regional associations, however, nine remained significant after conditioning on other lead variants within the *ABO* locus, including risk an increasing effect on anemias, for which rs550057 lead the genome-wide significant association signal (p = 4.7×10^-8^), visual field disturbances (p < 6.5×10^-5^) and diseases of the ear and mastoid process (p = 4.8×10^-5^). Replication of only two of the nine associations (other anemias and visual field defects) could be attempted in the UKBB due to poor phenotype matching and did not replicate; however, bearing relevance to the genome-wide significant finding in anemia, rs550057 led the association with red blood cell count in the UKBB (p = 1.3×10^-212^).^52^

#### Driver traits

IL-4 was the only driver trait of the multivariate association and has been implicated in the pathogenesis of many of the diseases associated with the locus. Aplastic anemia is considered to result primarily from immune-mediated bone marrow failure and an imbalance in Type I versus Type II T-cells that secrete IL-4 among other cytokines has been reported^53^. In endometriosis, IL-4 levels have been shown to be upregulated and induce the proliferation of endometriotic stromal cells^54, 55^.

#### eQTLs and pQTLs

The lead variant rs550057 is a pQTL impacting the levels of four proteins: ALPI, CHST15, FAM177A1 and JAG1 ^6^. Two of these proteins, carbohydrate sulfotransferase 15 (CHST15) and Jagged1 (JAG1), have been implicated in the pathogenesis of diseases associated with the locus. A small-interfering RNA targeting CHST15 improved myocardial function as well as reduced cardiac fibrosis, hypertrophy and secretion of proinflammatory cytokines in rats with chronic heart failure^56^. Upregulation of JAG1 has been reported in the endometrium of patients with endometriosis compared to controls^57^. Alagille Syndrome mainly caused by mutations in the JAG1 gene, is accompanied by congenital heart defects and varying degrees of hypercholesterolemia^58^

### *F5* gene locus

#### Multivariate association

An association with the *F5* locus was only detected by multivariate analysis (minimum univariate p = 1.1×10^-3^ for the lead variant from multivariate analysis) and the locus had not been previously associated with any of the biomarkers. The locus included two putative causal variants, rs3820060 and rs9332701, out of which the former was the lead variant from multivariate analysis (p = 6.15×10^−20^).

#### FinnGen disease associations

Three genome-wide significant missense variants in the *F5* locus (rs4524, rs4525, rs6032), all in high LD with one another (r^2^ > 0.98), were associated with nine diseases in FinnGen with four of these associations remaining significant after conditioning. Three of the four associations were protective for venous thromboembolism (VTE)-related endpoints (p < 6.9×10^-5^) and one increased the risk of fluid and electrolyte balance disruption, more specifically hypo-osmolality and hyponatraemia (p = 9.5×10^-5^, Supplementary Table 2). We replicated a previously reported protective effect of the missense variant rs4524 on VTE^59^ that remained significant after conditioning on factor V Leiden (rs6025; p = 1.5×10^-11^), a missense variant with a well-known risk-increasing effect on VTE^60^, while the hypo-osmolality and hyponatraemia association was novel. The VTE-related associations were replicated in the UKBB. A fourth missense variant in the locus (rs6027) increased the risk of four VTE-related diseases (p < 2.4×10^-5^), all of which remained significant after conditioning on the variant with the strongest association in the locus. These associations were not replicated in the UKBB.

#### Driver traits

The multivariate association in this locus had two driver traits: IL-4 and IL-12 both of which are relevant for coagulation as IL-12 has been shown to activate coagulation^61^ and cross-talk between the inflammatory and coagulation systems is extensive^62^.

#### eQTLs and pQTLs

The rs3820060 variant was an eQTL for the *F5* and *NME7* genes and was in the same LD-block (r^2^> 0.6) as many pQTLs affecting SEC13, NPTX2, SIG11, CAMK1, and TFPI levels. This block also included the three highly-correlated genome-wide significant missense variants mentioned above. Tissue factor pathway inhibitor (TFPI) is a major antithrombotic protein that inhibits thrombin and the external coagulation pathway. Low levels of TFPI increase the risk of venous thrombosis^63^ and TFPI has been shown to interact with the two driver traits IL-4 and IL-12^64, 65^. The other causal variant rs9332701 was a pQTL for F5^5^ and was in high LD (r^2^ = 0.97) with an eQTL for *NME7* and a pQTL for EHBP1^6^.

## DISCUSSION

We developed a novel method for multivariate GWAS follow-up analyses and demonstrated the considerable boost in power provided by multivariate GWAS using 12 highly correlated inflammatory markers. In total, four out of 11 genome-wide significant loci were detected only by multivariate analysis when adjusting univariate GWAS for multiple testing. At several loci, multivariate analysis also seemed to highlight more plausible candidates for causal variants than the univariate analyses. For example, in the *C1QA* locus, the lead variant in the univariate GWAS of the driver trait TNF-β was an intronic variant in the *EPHB2* gene, whereas the lead variant for the locus in the multivariate analysis was a Finnish-enriched missense variant located in the *C1QA* gene which has been previously associated with immunologic diseases. Our multivariate analysis may point towards a plausible mechanism underlying these associations via TNF-β levels.

Although both univariate and multivariate scans have previously been applied to these biomarkers^1, 66^, these studies have suffered from the lack of essential follow-up analyses due to the absence of beta estimates in multivariate summary statistics. Our novel method enables two key follow-up analyses for multivariate GWAS: fine-mapping and trait prioritization. Our method solves the problem of missing effect sizes and standard errors required for fine-mapping by an extension of metaCCA followed by LCP-GWAS. This process allows for the transformation of CCA-based multivariate GWAS results into univariate summary statistics and thus extends the use of FINEMAP and other summary statistics-based tools to multivariate GWAS. Fine-mapping complex multivariate associations allows for assessing causality of the variants within the associated loci. This has not been previously feasible. We also further describe the multivariate associations by determining the traits driving the associations using MetaPhat. This workflow allows the identification of both the variants and traits underlying the multivariate associations.

Our study also elucidates the advantage of multivariate analysis combined with large biobank-based phenome-wide screening by discovering multiple novel disease associations. For example, in the *GP6* locus we observe a novel risk-increasing association between the Finnish enriched rare missense variant rs199588110 and benign neoplasms of meninges. Altogether, a majority of the observed disease associations were for variants in the *F5* and *ABO* loci that were only detected by multivariate GWAS. All these associations, including a genome-wide significant association with anemia that replicated in the UKBB as an effect on red blood cell count, would have gone undetected had we used univariate GWAS. In addition to disease association discovery, our workflow promotes increasing insight into the pathophysiology underlying the associations by identifying the biomarkers driving the associations. Detailed exploration of biological evidence including eQTLs and pQTLs in the *GP6, SERPINE2, ABO,* and *F5* loci orthogonally supports our evidence of causal variants and driver traits. For example, in the *SERPINE2* locus one of the three putative causal variants rs13412535 increased the risk of hypertrophic skin disorders in FinnGen and was a pQTL for PDGF-BB ^6^ that is considered a key player in the pathogenesis of hypertrophic scars ^50^, increasing evidence of the biologically relevant functions of this variant.

These methodological development and novel findings notwithstanding, our study has some limitations. First, our newly developed workflow for multivariate fine-mapping requires individual level genotype and phenotype data, problematic for some analysis settings. Additionally, the LCPs are optimized for the lead variants, potentially resulting in overestimation of the causal probability of these variants. We did not, however, see evidence of this in the *F5* locus where we constructed LCPs for two missense variants in addition to the lead variant with no significant changes in the causal probabilities of the variants. We also acknowledge that the credible sets we chose for follow-up may not encompass all causal signals within the multivariate associations. The credible sets excluded due to low LD may arise from multiple signals included in the same set, resulting in small LD within the set. Further, some disease associations require replication and follow-up analyses.

On the other hand, our study has many strengths. First, a prospective cohort study was used to assess deep phenotype data rarely available at large scale. Second, we are the first to present phenome-wide results from FinnGen, a very large and well-phenotyped Finnish biobank study, and also make use of the UKBB, in disease association follow-up, ensuring enough power for disease association detection. Finland has a public healthcare system and national health registries, which enable the vast and accurate phenotyping in FinnGen. Besides FinnGen, an additional advantage to performing the study in Finns is that deleterious variants are enriched in the Finnish population due to population history^21^. Furthermore, our reference panel for genotype imputation is from the same population as our discovery and follow-up data sets, which, as demonstrated also by others^67, 68^, allows us to study variants that are enriched (and often unique) in the study-specific population.

In conclusion, we developed a novel workflow for multivariate GWAS discovery and follow-up analyses, including fine-mapping and identification of driver traits, and thus promote the advancement of powerful multivariate methods in genomic analyses. We demonstrate the benefit of applying this workflow by identifying novel associations and further describing previously reported associations with both biomarkers and diseases using a set of inflammatory markers. We show that compared to univariate analyses, multivariate analysis of biomarker data combined with large biobank-based PheWAS reveals a considerably increased number of novel genetic associations with several diseases.

## AKNOWLEDGMENTS

We would like to thank Lea Urpa for proofreading, and Sari Kivikko, Huei-Yi Shen and Ulla Tuomainen for management assistance. We would like to thank all participants of the FINRISK, FinnGen and UKBB studies for their generous participation. The FINRISK data used for the research were obtained from THL Biobank. This research has been conducted using the UK Biobank Resource with application number 22627.

This work was supported by the Academy of Finland Center of Excellence in Complex Disease Genetics [Grant No 312062 to S.R., 312074 to A.P., 312075 to M.D.]; Academy of Finland [Grant No 285380 to S.R, 128650 to A.P.]; the Finnish Foundation for Cardiovascular Research [to S.R., V.S., and A.P.]; the Sigrid Jusélius Foundation [to S.R. and A.P.]; University of Helsinki HiLIFE Fellow grants 2017-2020 [to S.R.]; Foundation and the Horizon 2020 Research and Innovation Programme [grant number 667301 (COSYN) to A.P.]; the Doctoral Programme in Population Health, University of Helsinki [to J.J.P. and S.E.R.]; and The Finnish Medical Foundation [to J.J.P.]. The FinnGen project is funded by two grants from Business Finland (HUS 4685/31/2016 and UH 4386/31/2016) and nine industry partners (AbbVie, AstraZeneca, Biogen, Celgene, Genentech, GSK, MSD, Pfizer and Sanofi). Following biobanks are acknowledged for collecting the FinnGen project samples: Auria Biobank (https://www.auria.fi/biopankki/en), THL Biobank (https://thl.fi/fi/web/thl-biopankki), Helsinki Biobank (https://www.terveyskyla.fi/helsinginbiopankki/en), Northern Finland Biobank Borealis (https://www.ppshp.fi/Tutkimus-ja-opetus/Biopankki), Finnish Clinical Biobank Tampere (https://www.tays.fi/en-US/Research_and_development/Finnish_Clinical_Biobank_Tampere), Biobank of Eastern Finland (https://ita-suomenbiopankki.fi/), Central Finland Biobank (https://www.ksshp.fi/fi-FI/Potilaalle/Biopankki), Finnish Red Cross Blood Service Biobank (https://www.bloodservice.fi/Research%20Projects/biobanking).

The funders had no role in study design, data collection and analysis, decision to publish, or preparation of the manuscript.

## Contributors of FinnGen

### Steering Committee

Aarno Palotie Institute for Molecular Medicine Finland, HiLIFE, University of Helsinki, Finland

Mark Daly Institute for Molecular Medicine Finland, HiLIFE, University of Helsinki, Finland

#### Pharmaceutical companies

Howard Jacob Abbvie, Chicago, IL, United States

Athena Matakidou Astra Zeneca, Cambridge, United Kingdom

Heiko Runz Biogen, Cambridge, MA, United States

Sally John Biogen, Cambridge, MA, United States

Robert Plenge Celgene, Summit, NJ, United States

Mark McCarthy Genentech, San Francisco, CA, United States

Julie Hunkapiller Genentech, San Francisco, CA, United States

Meg Ehm GlaxoSmithKline, Brentford, United Kingdom

Dawn Waterworth GlaxoSmithKline, Brentford, United Kingdom

Caroline Fox Merck, Kenilworth, NJ, United States

Anders Malarstig Pfizer, New York, NY, United States

Kathy Klinger Sanofi, Paris, France

Kathy Call Sanofi, Paris, France

#### University of Helsinki & Biobanks

Tomi Mäkelä HiLIFE, University of Helsinki, Finland, Finland

Jaakko Kaprio Institute for Molecular Medicine Finland, HiLIFE, Helsinki, Finland, Finland

Petri Virolainen Auria Biobank / Univ. of Turku / Hospital District of Southwest Finland, Turku, Finland

Kari Pulkki Auria Biobank / Univ. of Turku / Hospital District of Southwest Finland, Turku, Finland

Terhi Kilpi THL Biobank / The National Institute of Health and Welfare Helsinki, Finland

Markus Perola THL Biobank / The National Institute of Health and Welfare Helsinki, Finland

Jukka Partanen Finnish Red Cross Blood Service / Finnish Hematology Registry and Clinical Biobank, Helsinki, Finland

Anne Pitkäranta Hospital District of Helsinki and Uusimaa, Helsinki, Finland

Riitta Kaarteenaho Northern Finland Biobank Borealis / University of Oulu / Northern Ostrobothnia Hospital District, Oulu, Finland

Seppo Vainio Northern Finland Biobank Borealis / University of Oulu / Northern Ostrobothnia Hospital District, Oulu, Finland

Kimmo Savinainen Finnish Clinical Biobank Tampere **/** University of Tampere / Pirkanmaa Hospital District, Tampere, Finland

Veli-Matti Kosma Biobank of Eastern Finland / University of Eastern Finland / Northern Savo Hospital District, Kuopio, Finland

Urho Kujala Central Finland Biobank / University of Jyväskylä / Central Finland Health Care District, Jyväskylä, Finland

#### Other Experts/ Non-Voting Members

Outi Tuovila Business Finland, Helsinki, Finland

Minna Hendolin Business Finland, Helsinki, Finland

Raimo Pakkanen Business Finland, Helsinki, Finland

### Scientific Committee

#### Pharmaceutical companies

Jeff Waring Abbvie, Chicago, IL, United States

Bridget Riley-Gillis Abbvie, Chicago, IL, United States

Athena Matakidou Astra Zeneca, Cambridge, United Kingdom

Heiko Runz Biogen, Cambridge, MA, United States

Jimmy Liu Biogen, Cambridge, MA, United States

Shameek Biswas Celgene, Summit, NJ, United States

Julie Hunkapiller Genentech, San Francisco, CA, United States

Dawn Waterworth GlaxoSmithKline, Brentford, United Kingdom

Meg Ehm GlaxoSmithKline, Brentford, United Kingdom

Dorothee Diogo Merck, Kenilworth, NJ, United States

Caroline Fox Merck, Kenilworth, NJ, United States

Anders Malarstig Pfizer, New York, NY, United States

Catherine Marshall Pfizer, New York, NY, United States

Xinli Hu Pfizer, New York, NY, United States

Kathy Call Sanofi, Paris, France

Kathy Klinger Sanofi, Paris, France

Matthias Gossel Sanofi, Paris, France

#### University of Helsinki & Biobanks

Samuli Ripatti Institute for Molecular Medicine Finland, HiLIFE, Helsinki, Finland

Johanna Schleutker Auria Biobank / Univ. of Turku / Hospital District of Southwest Finland, Turku, Finland

Markus Perola THL Biobank / The National Institute of Health and Welfare Helsinki, Finland

Mikko Arvas Finnish Red Cross Blood Service / Finnish Hematology Registry and Clinical Biobank, Helsinki, Finland

Olli Carpen Hospital District of Helsinki and Uusimaa, Helsinki, Finland

Reetta Hinttala Northern Finland Biobank Borealis / University of Oulu / Northern Ostrobothnia Hospital District, Oulu, Finland

Johannes Kettunen Northern Finland Biobank Borealis / University of Oulu / Northern Ostrobothnia Hospital District, Oulu, Finland

Reijo Laaksonen Finnish Clinical Biobank Tampere / University of Tampere / Pirkanmaa Hospital District, Tampere, Finland

Arto Mannermaa Biobank of Eastern Finland / University of Eastern Finland / Northern Savo Hospital District, Kuopio, Finland

Juha Paloneva Central Finland Biobank / University of Jyväskylä / Central Finland Health Care District, Jyväskylä, Finland

Urho Kujala Central Finland Biobank / University of Jyväskylä / Central Finland Health Care District, Jyväskylä, Finland

#### Other Experts/ Non-Voting Members

Outi Tuovila Business Finland, Helsinki, Finland

Minna Hendolin Business Finland, Helsinki, Finland

Raimo Pakkanen Business Finland, Helsinki, Finland

### Clinical Groups

#### Neurology Group

Hilkka Soininen Northern Savo Hospital District, Kuopio, Finland

Valtteri Julkunen Northern Savo Hospital District, Kuopio, Finland

Anne Remes Northern Ostrobothnia Hospital District, Oulu, Finland

Reetta Kälviäinen Northern Savo Hospital District, Kuopio, Finland

Mikko Hiltunen Northern Savo Hospital District, Kuopio, Finland

Jukka Peltola Pirkanmaa Hospital District, Tampere, Finland

Pentti Tienari Hospital District of Helsinki and Uusimaa, Helsinki, Finland

Juha Rinne Hospital District of Southwest Finland, Turku, Finland

Adam Ziemann Abbvie, Chicago, IL, United States

Jeffrey Waring Abbvie, Chicago, IL, United States

Sahar Esmaeeli Abbvie, Chicago, IL, United States

Nizar Smaoui Abbvie, Chicago, IL, United States

Anne Lehtonen Abbvie, Chicago, IL, United States

Susan Eaton Biogen, Cambridge, MA, United States

Heiko Runz Biogen, Cambridge, MA, United States

Sanni Lahdenperä Biogen, Cambridge, MA, United States

Shameek Biswas Celgene, Summit, NJ, United States

John Michon Genentech, San Francisco, CA, United States

Geoff Kerchner Genentech, San Francisco, CA, United States

Julie Hunkapiller Genentech, San Francisco, CA, United States

Natalie Bowers Genentech, San Francisco, CA, United States

Edmond Teng Genentech, San Francisco, CA, United States

John Eicher Merck, Kenilworth, NJ, United States

Vinay Mehta Merck, Kenilworth, NJ, United States

Padhraig Gormley Merck, Kenilworth, NJ, United States

Kari Linden Pfizer, New York, NY, United States

Christopher Whelan Pfizer, New York, NY, United States

Fanli Xu GlaxoSmithKline, Brentford, United Kingdom

David Pulford GlaxoSmithKline, Brentford, United Kingdom

#### Gastroenterology Group

Martti Färkkilä Hospital District of Helsinki and Uusimaa, Helsinki, Finland

Sampsa Pikkarainen Hospital District of Helsinki and Uusimaa, Helsinki, Finland

Airi Jussila Pirkanmaa Hospital District, Tampere, Finland

Timo Blomster Northern Ostrobothnia Hospital District, Oulu, Finland

Mikko Kiviniemi Northern Savo Hospital District, Kuopio, Finland

Markku Voutilainen Hospital District of Southwest Finland, Turku, Finland

Bob Georgantas Abbvie, Chicago, IL, United States

Graham Heap Abbvie, Chicago, IL, United States

Jeffrey Waring Abbvie, Chicago, IL, United States

Nizar Smaoui Abbvie, Chicago, IL, United States

Fedik Rahimov Abbvie, Chicago, IL, United States

Anne Lehtonen Abbvie, Chicago, IL, United States

Keith Usiskin Celgene, Summit, NJ, United States

Joseph Maranville Celgene, Summit, NJ, United States

Tim Lu Genentech, San Francisco, CA, United States

Natalie Bowers Genentech, San Francisco, CA, United States

Danny Oh Genentech, San Francisco, CA, United States

John Michon Genentech, San Francisco, CA, United States

Vinay Mehta Merck, Kenilworth, NJ, United States

Kirsi Kalpala Pfizer, New York, NY, United States

Melissa Miller Pfizer, New York, NY, United States

Xinli Hu Pfizer, New York, NY, United States

Linda McCarthy GlaxoSmithKline, Brentford, United Kingdom

#### Rheumatology Group

Kari Eklund Hospital District of Helsinki and Uusimaa, Helsinki, Finland

Antti Palomäki Hospital District of Southwest Finland, Turku, Finland

Pia Isomäki Pirkanmaa Hospital District, Tampere, Finland

Laura Pirilä Hospital District of Southwest Finland, Turku, Finland

Oili Kaipiainen-Seppänen Northern Savo Hospital District, Kuopio, Finland

Johanna Huhtakangas Northern Ostrobothnia Hospital District, Oulu, Finland

Bob Georgantas Abbvie, Chicago, IL, United States

Jeffrey Waring Abbvie, Chicago, IL, United States

Fedik Rahimov Abbvie, Chicago, IL, United States

Apinya Lertratanakul Abbvie, Chicago, IL, United States

Nizar Smaoui Abbvie, Chicago, IL, United States

Anne Lehtonen Abbvie, Chicago, IL, United States

David Close Astra Zeneca, Cambridge, United Kingdom

Marla Hochfeld Celgene, Summit, NJ, United States

Natalie Bowers Genentech, San Francisco, CA, United States

John Michon Genentech, San Francisco, CA, United States

Dorothee Diogo Merck, Kenilworth, NJ, United States

Vinay Mehta Merck, Kenilworth, NJ, United States

Kirsi Kalpala Pfizer, New York, NY, United States

Nan Bing Pfizer, New York, NY, United States

Xinli Hu Pfizer, New York, NY, United States

Jorge Esparza Gordillo GlaxoSmithKline, Brentford, United Kingdom

Nina Mars Institute for Molecular Medicine Finland, HiLIFE, Helsinki, Finland

#### Pulmonology Group

Tarja Laitinen Pirkanmaa Hospital District, Tampere, Finland

Margit Pelkonen Northern Savo Hospital District, Kuopio, Finland

Paula Kauppi Hospital District of Helsinki and Uusimaa, Helsinki, Finland

Hannu Kankaanranta Pirkanmaa Hospital District, Tampere, Finland

Terttu Harju Northern Ostrobothnia Hospital District, Oulu, Finland

Nizar Smaoui Abbvie, Chicago, IL, United States

David Close Astra Zeneca, Cambridge, United Kingdom

Steven Greenberg Celgene, Summit, NJ, United States

Hubert Chen Genentech, San Francisco, CA, United States

Natalie Bowers Genentech, San Francisco, CA, United States

John Michon Genentech, San Francisco, CA, United States

Vinay Mehta Merck, Kenilworth, NJ, United States

Jo Betts GlaxoSmithKline, Brentford, United Kingdom

Soumitra Ghosh GlaxoSmithKline, Brentford, United Kingdom

#### Cardiometabolic Diseases Group

Veikko Salomaa The National Institute of Health and Welfare Helsinki, Finland

Teemu Niiranen The National Institute of Health and Welfare Helsinki, Finland

Markus Juonala Hospital District of Southwest Finland, Turku, Finland

Kaj Metsärinne Hospital District of Southwest Finland, Turku, Finland

Mika Kähönen Pirkanmaa Hospital District, Tampere, Finland

Juhani Junttila Northern Ostrobothnia Hospital District, Oulu, Finland

Markku Laakso Northern Savo Hospital District, Kuopio, Finland

Jussi Pihlajamäki Northern Savo Hospital District, Kuopio, Finland

Juha Sinisalo Hospital District of Helsinki and Uusimaa, Helsinki, Finland

Marja-Riitta Taskinen Hospital District of Helsinki and Uusimaa, Helsinki, Finland

Tiinamaija Tuomi Hospital District of Helsinki and Uusimaa, Helsinki, Finland

Jari Laukkanen Central Finland

Health Care District, Jyväskylä, Finland

Ben Challis Astra Zeneca, Cambridge, United Kingdom

Andrew Peterson Genentech, San Francisco, CA, United States

Julie Hunkapiller Genentech, San Francisco, CA, United States

Natalie Bowers Genentech, San Francisco, CA, United States

John Michon Genentech, San Francisco, CA, United States

Dorothee Diogo Merck, Kenilworth, NJ, United States

Audrey Chu Merck, Kenilworth, NJ, United States

Vinay Mehta Merck, Kenilworth, NJ, United States

Jaakko Parkkinen Pfizer, New York, NY, United States

Melissa Miller Pfizer, New York, NY, United States

Anthony Muslin Sanofi, Paris, France

Dawn Waterworth GlaxoSmithKline, Brentford, United Kingdom

#### Oncology Group

Heikki Joensuu Hospital District of Helsinki and Uusimaa, Helsinki, Finland

Tuomo Meretoja Hospital District of Helsinki and Uusimaa, Helsinki, Finland

Olli Carpen Hospital District of Helsinki and Uusimaa, Helsinki, Finland

Lauri Aaltonen Hospital District of Helsinki and Uusimaa, Helsinki, Finland

Annika Auranen Pirkanmaa Hospital District, Tampere, Finland

Peeter Karihtala Northern Ostrobothnia Hospital District, Oulu, Finland

Saila Kauppila Northern Ostrobothnia Hospital District, Oulu, Finland

Päivi Auvinen Northern Savo Hospital District, Kuopio, Finland

Klaus Elenius Hospital District of Southwest Finland, Turku, Finland

Relja Popovic Abbvie, Chicago, IL, United States

Jeffrey Waring Abbvie, Chicago, IL, United States

Bridget Riley-Gillis Abbvie, Chicago, IL, United States

Anne Lehtonen Abbvie, Chicago, IL, United States

Athena Matakidou Astra Zeneca, Cambridge, United Kingdom

Jennifer Schutzman Genentech, San Francisco, CA, United States

Julie Hunkapiller Genentech, San Francisco, CA, United States

Natalie Bowers Genentech, San Francisco, CA, United States

John Michon Genentech, San Francisco, CA, United States

Vinay Mehta Merck, Kenilworth, NJ, United States

Andrey Loboda Merck, Kenilworth, NJ, United States

Aparna Chhibber Merck, Kenilworth, NJ, United States

Heli Lehtonen Pfizer, New York, NY, United States

Stefan McDonough Pfizer, New York, NY, United States

Marika Crohns Sanofi, Paris, France

Diptee Kulkarni GlaxoSmithKline, Brentford, United Kingdom

#### Opthalmology Group

Kai Kaarniranta Northern Savo Hospital District, Kuopio, Finland

Joni Turunen Hospital District of Helsinki and Uusimaa, Helsinki, Finland

Terhi Ollila Hospital District of Helsinki and Uusimaa, Helsinki, Finland

Sanna Seitsonen Hospital District of Helsinki and Uusimaa, Helsinki, Finland

Hannu Uusitalo Pirkanmaa Hospital District, Tampere, Finland

Vesa Aaltonen Hospital District of Southwest Finland, Turku, Finland

Hannele Uusitalo-Järvinen Pirkanmaa Hospital District, Tampere, Finland

Marja Luodonpää Northern Ostrobothnia Hospital District, Oulu, Finland

Nina Hautala Northern Ostrobothnia Hospital District, Oulu, Finland

Heiko Runz Biogen, Cambridge, MA, United States

Erich Strauss Genentech, San Francisco, CA, United States

Natalie Bowers Genentech, San Francisco, CA, United States

Hao Chen Genentech, San Francisco, CA, United States

John Michon Genentech, San Francisco, CA, United States

Anna Podgornaia Merck, Kenilworth, NJ, United States

Vinay Mehta Merck, Kenilworth, NJ, United States

Dorothee Diogo Merck, Kenilworth, NJ, United States

Joshua Hoffman GlaxoSmithKline, Brentford, United Kingdom

#### Dermatology Group

Kaisa Tasanen Northern Ostrobothnia Hospital District, Oulu, Finland

Laura Huilaja Northern Ostrobothnia Hospital District, Oulu, Finland

Katariina Hannula-Jouppi Hospital District of Helsinki and Uusimaa, Helsinki, Finland

Teea Salmi Pirkanmaa Hospital District, Tampere, Finland

Sirkku Peltonen Hospital District of Southwest Finland, Turku, Finland

Leena Koulu Hospital District of Southwest Finland, Turku, Finland

Ilkka Harvima Northern Savo Hospital District, Kuopio, Finland

Kirsi Kalpala Pfizer, New York, NY, United States

Ying Wu Pfizer, New York, NY, United States

David Choy Genentech, San Francisco, CA, United States

John Michon Genentech, San Francisco, CA, United States

Nizar Smaoui Abbvie, Chicago, IL, United States

Fedik Rahimov Abbvie, Chicago, IL, United States

Anne Lehtonen Abbvie, Chicago, IL, United States

Dawn Waterworth GlaxoSmithKline, Brentford, United Kingdom

### FinnGen Teams

#### Administration Team

Anu Jalanko Institute for Molecular Medicine Finland, HiLIFE, University of Helsinki, Finland

Risto Kajanne Institute for Molecular Medicine Finland, HiLIFE, University of Helsinki, Finland

Ulrike Lyhs Institute for Molecular Medicine Finland, HiLIFE, University of Helsinki, Finland

#### Communication

Mari Kaunisto Institute for Molecular Medicine Finland, HiLIFE, University of Helsinki, Finland

#### Analysis Team

Justin Wade Davis Abbvie, Chicago, IL, United States

Bridget Riley-Gillis Abbvie, Chicago, IL, United States

Danjuma Quarless Abbvie, Chicago, IL, United States

Slavé Petrovski Astra Zeneca, Cambridge, United Kingdom

Jimmy Liu Biogen, Cambridge, MA, United States

Chia-Yen Chen Biogen, Cambridge, MA, United States

Paola Bronson Biogen, Cambridge, MA, United States

Robert Yang Celgene, Summit, NJ, United States

Joseph Maranville Celgene, Summit, NJ, United States

Shameek Biswas Celgene, Summit, NJ, United States

Diana Chang Genentech, San Francisco, CA, United States

Julie Hunkapiller Genentech, San Francisco, CA, United States

Tushar Bhangale Genentech, San Francisco, CA, United States

Natalie Bowers Genentech, San Francisco, CA, United States

Dorothee Diogo Merck, Kenilworth, NJ, United States

Emily Holzinger Merck, Kenilworth, NJ, United States

Padhraig Gormley Merck, Kenilworth, NJ, United States

Xulong Wang Merck, Kenilworth, NJ, United States

Xing Chen Pfizer, New York, NY, United States

Åsa Hedman Pfizer, New York, NY, United States

Kirsi Auro GlaxoSmithKline, Brentford, United Kingdom

Clarence Wang Sanofi, Paris, France

Ethan Xu Sanofi, Paris, France

Franck Auge Sanofi, Paris, France

Clement Chatelain Sanofi, Paris, France

Mitja Kurki Institute for Molecular Medicine Finland, HiLIFE, University of Helsinki, Finland / Broad Institute, Cambridge, MA, United States

Samuli Ripatti Institute for Molecular Medicine Finland, HiLIFE, University of Helsinki, Finland

Mark Daly Institute for Molecular Medicine Finland, HiLIFE, University of Helsinki, Finland

Juha Karjalainen Institute for Molecular Medicine Finland, HiLIFE, University of Helsinki, Finland / Broad Institute, Cambridge, MA, United States

Aki Havulinna Institute for Molecular Medicine Finland, HiLIFE, University of Helsinki, Finland

Anu Jalanko Institute for Molecular Medicine Finland, HiLIFE, University of Helsinki, Finland

Kimmo Palin University of Helsinki, Helsinki, Finland

Priit Palta Institute for Molecular Medicine Finland, HiLIFE, University of Helsinki, Finland

Pietro Della Briotta Parolo Institute for Molecular Medicine Finland, HiLIFE, University of Helsinki, Finland

Wei Zhou Broad Institute, Cambridge, MA, United States

Susanna Lemmelä Institute for Molecular Medicine Finland, HiLIFE, University of Helsinki, Finland

Manuel Rivas University of Stanford, Stanford, CA, United States

Jarmo Harju Institute for Molecular Medicine Finland, HiLIFE, University of Helsinki, Finland

Aarno Palotie Institute for Molecular Medicine Finland, HiLIFE, University of Helsinki, Finland

Arto Lehisto Institute for Molecular Medicine Finland, HiLIFE, University of Helsinki, Finland

Andrea Ganna Institute for Molecular Medicine Finland, HiLIFE, University of Helsinki, Finland

Vincent Llorens Institute for Molecular Medicine Finland, HiLIFE, University of Helsinki, Finland

Antti Karlsson Auria Biobank / Univ. of Turku / Hospital District of Southwest Finland, Turku, Finland

Kati Kristiansson THL Biobank / The National Institute of Health and Welfare Helsinki, Finland

Mikko Arvas Finnish Red Cross Blood Service / Finnish Hematology Registry and Clinical Biobank, Helsinki, Finland

Kati Hyvärinen Finnish Red Cross Blood Service / Finnish Hematology Registry and Clinical Biobank, Helsinki, Finland

Jarmo Ritari Finnish Red Cross Blood Service / Finnish Hematology Registry and Clinical Biobank, Helsinki, Finland

Tiina Wahlfors Finnish Red Cross Blood Service / Finnish Hematology Registry and Clinical Biobank, Helsinki, Finland

Miika Koskinen Hospital District of Helsinki and Uusimaa, Helsinki, Finland BB/HUS/Univ Hosp Districts

Olli Carpen Hospital District of Helsinki and Uusimaa, Helsinki, Finland BB/HUS/Univ Hosp Districts

Johannes Kettunen Northern Finland Biobank Borealis / University of Oulu / Northern Ostrobothnia Hospital District, Oulu, Finland

Katri Pylkäs Northern Finland Biobank Borealis / University of Oulu / Northern Ostrobothnia Hospital District, Oulu, Finland

Marita Kalaoja Northern Finland Biobank Borealis / University of Oulu / Northern Ostrobothnia Hospital District, Oulu, Finland

Minna Karjalainen Northern Finland Biobank Borealis / University of Oulu / Northern Ostrobothnia Hospital District, Oulu, Finland

Tuomo Mantere Northern Finland Biobank Borealis / University of Oulu / Northern Ostrobothnia Hospital District, Oulu, Finland

Eeva Kangasniemi Finnish Clinical Biobank Tampere / University of Tampere / Pirkanmaa Hospital District, Tampere, Finland

Sami Heikkinen Biobank of Eastern Finland / University of Eastern Finland / Northern Savo Hospital District, Kuopio, Finland

Arto Mannermaa Biobank of Eastern Finland / University of Eastern Finland / Northern Savo Hospital District, Kuopio, Finland

Eija Laakkonen Central Finland Biobank / University of Jyväskylä / Central Finland Health Care District, Jyväskylä, Finland

Juha Kononen Central Finland Biobank / University of Jyväskylä / Central Finland Health Care District, Jyväskylä, Finland

#### Sample Collection Coordination

Anu Loukola Hospital District of Helsinki and Uusimaa, Helsinki, Finland

#### Sample Logistics

Päivi Laiho THL Biobank / The National Institute of Health and Welfare Helsinki, Finland

Tuuli Sistonen THL Biobank / The National Institute of Health and Welfare Helsinki, Finland

Essi Kaiharju THL Biobank / The National Institute of Health and Welfare Helsinki, Finland

Markku Laukkanen THL Biobank / The National Institute of Health and Welfare Helsinki, Finland

Elina Järvensivu THL Biobank / The National Institute of Health and Welfare Helsinki, Finland

Sini Lähteenmäki THL Biobank / The National Institute of Health and Welfare Helsinki, Finland

Lotta Männikkö THL Biobank / The National Institute of Health and Welfare Helsinki, Finland

Regis Wong THL Biobank / The National Institute of Health and Welfare Helsinki, Finland

#### Registry Data Operations

Kati Kristiansson THL Biobank / The National Institute of Health and Welfare Helsinki, Finland

Hannele Mattsson THL Biobank / The National Institute of Health and Welfare Helsinki, Finland

Susanna Lemmelä Institute for Molecular Medicine Finland, HiLIFE, University of Helsinki, Finland

Tero Hiekkalinna THL Biobank / The National Institute of Health and Welfare Helsinki, Finland

Manuel González Jiménez THL Biobank / The National Institute of Health and Welfare Helsinki, Finland

#### Genotyping

Kati Donner Institute for Molecular Medicine Finland, HiLIFE, University of Helsinki, Finland

#### Sequencing Informatics

Priit Palta Institute for Molecular Medicine Finland, HiLIFE, University of Helsinki, Finland

Kalle Pärn Institute for Molecular Medicine Finland, HiLIFE, University of Helsinki, Finland

Javier Nunez-Fontarnau Institute for Molecular Medicine Finland, HiLIFE, University of Helsinki, Finland

#### Data Management and IT Infrastructure

Jarmo Harju Institute for Molecular Medicine Finland, HiLIFE, University of Helsinki, Finland

Elina Kilpeläinen Institute for Molecular Medicine Finland, HiLIFE, University of Helsinki, Finland

Timo P. Sipilä Institute for Molecular Medicine Finland, HiLIFE, University of Helsinki, Finland

Georg Brein Institute for Molecular Medicine Finland, HiLIFE, University of Helsinki, Finland

Alexander Dada Institute for Molecular Medicine Finland, HiLIFE, University of Helsinki, Finland

Ghazal Awaisa Institute for Molecular Medicine Finland, HiLIFE, University of Helsinki, Finland

Anastasia Shcherban Institute for Molecular Medicine Finland, HiLIFE, University of Helsinki, Finland

Tuomas Sipilä Institute for Molecular Medicine Finland, HiLIFE, University of Helsinki, Finland

#### Clinical Endpoint Development

Hannele Laivuori Institute for Molecular Medicine Finland, HiLIFE, University of Helsinki, Finland

Aki Havulinna Institute for Molecular Medicine Finland, HiLIFE, University of Helsinki, Finland

Susanna Lemmelä Institute for Molecular Medicine Finland, HiLIFE, University of Helsinki, Finland

Tuomo Kiiskinen Institute for Molecular Medicine Finland, HiLIFE, University of Helsinki, Finland

#### Trajectory Team

Tarja Laitinen Pirkanmaa Hospital District, Tampere, Finland

Harri Siirtola University of Tampere, Tampere, Finland

Javier Gracia Tabuenca University of Tampere, Tampere, Finland

#### Biobank Directors

Lila Kallio Auria Biobank, Turku, Finland

Sirpa Soini THL Biobank, Helsinki, Finland

Jukka Partanen Blood Service Biobank, Helsinki, Finland

Kimmo Pitkänen Helsinki Biobank, Helsinki, Finland

Seppo Vainio Northern Finland Biobank Borealis, Oulu, Finland

Kimmo Savinainen Tampere Biobank, Tampere, Finland

Veli-Matti Kosma Biobank of Eastern Finland, Kuopio, Finland

Teijo Kuopio Central Finland Biobank, Jyväskylä, Finland

