## Supplementary Material for "An expanded analysis framework for multivariate GWAS connects inflammatory biomarkers to functional variants and disease"

#### METHODS

##### Hierarchical clustering of quantitative traits

Clusters of highly correlated quantitative traits within 66 quantitative traits from the FINRISK study (collection years 1992-2012, sample sizes ranging from 4,792 to 26,717) were identified using hierarchical clustering. The pairwise Pearson correlation coefficients were calculated amongst inverse-rank normalized age and sex adjusted residuals of the 66 traits using complete samples for the two traits. The correlation structure of the traits is shown in Supplementary Figure 2. Hierarchical clustering was performed using the Ward agglomeration method and one minus the absolute correlation coefficient as the dissimilarity metric. P-values for the clusters were obtained via multiscale bootstrap resampling[1] . Clusters with over three traits and p-values passing the significance level 0.01 were considered.

##### Chip genotype data processing and QC

Samples were genotyped with Illumina (Illumina Inc., San Diego, CA, USA) and Affymetrix arrays (Thermo Fisher Scientific, Santa Clara, CA, USA). Genotype calls were made with GenCall and zCall algorithms for Illumina and AxiomGT1 algorithm for Affymetrix data. Chip genotyping data produced with previous chip platforms and reference genome builds were lifted over to build version 38 (GRCh38/hg38) following the protocol described her: dx.doi.org/10.17504/protocols.io.nqtddwn. In sample-wise quality control, individuals with ambiguous gender, high genotype missingness (>5%), excess heterozygosity (±4SD) and non-Finnish ancestry were removed. In variant-wise quality control variants with high missingness (>2%), low HWE p-value ($<1\times10^{-6}$) and minor allele count, MAC<3 were removed.

**Haplotype phasing and genotype imputation**

Chip-genotyped samples were pre-phased with Eagle 2.3.5 (https://data.broadinstitute.org/alkesgroup/Eagle/) with the default parameters, except the number of conditioning haplotypes was set to 20,000. Genotype imputation was carried out by using the population-specific SISu v3 imputation reference panel with Beagle 4.1 (version08Jun17.d8b, https://faculty.washington.edu/browning/beagle/b4_1.html) as described here: dx.doi.org/10.17504/protocols.io.nmndc5e. Post-imputation variant-wise quality-control involved checking expected conformity of the imputation INFO-values distribution and MAF differences between the target dataset and the imputation reference panel.

**Post-imputation QC**

Genotype imputation was followed by post-imputation sample QC, wherein 340 duplicate samples (kinship > 0.45) and 102 additional samples were excluded based on singleton count > 20, number of heterozygous calls falling out of hard called bounds (lower-bound 20,700, upper-bound 22,450), and over 4 standard deviation difference from the mean in number of insertion alternate alleles, number of deletion alternate alleles, insertion/deletion allele ratio, transition/transversion ratio and number of SNP alternate alleles. Sample QC was performed using 74,506 variants passing strict variant QC (autosomal single nucleotide polymorphisms with imputation INFO score > 0.99, 0.05 ≤ AF ≤ 0.95, HWE p-value > 0.001, call rate > 0.99, linkage disequilibrium (LD)-pruned with $r^2$ threshold 0.1 and 1 MB window-size), where possible (number of heterozygous calls, number of SNP alternate alleles, transition/transversion ratio). When the strict variant set could not be used (singletons, insertions, deletions, insertion/deletion ratio), variants were restricted to those with imputation INFO score ≥ 0.7. Lastly, genetic variants with imputation INFO score < 0.8, minor allele frequency < 0.002 or HWE p-value < $1\times10^{-6}$ were removed.

**Principal component analysis**

Principal component (PC) analysis for 26,717 FINIRISK individuals was performed using 73,072 independent high-quality variants (autosomal single nucleotide polymorphisms with imputation info score > 0.99, 0.05 ≤ AF ≤ 0.95, HWE p-value > 0.001, call rate > 0.99, excluding high-LD regions, LD-pruned with $r^2$ threshold 0.1 and 1 MB window-size). For PC analysis 2,119 related (kinship > 0.1) were excluded. First 10 PCs were calculated for 24,598 unrelated individuals, and SNP weights were extracted. Using those weights, first 10 PCs were projected for the 2,119 individuals excluded due to relatedness.

**FinnGen ethical statements**

For the Finnish Institute of Health and Welfare (THL) driven FinnGen preparatory project (here called FinnGen), all patients and control subjects had provided informed consent for biobank research, based on the Finnish Biobank Act. Alternatively, older cohorts were based on study specific consents and later transferred to the THL Biobank after approval by Valvira, the National Supervisory Authority for Welfare and Health. Recruitment protocols followed the biobank protocols approved by Valvira. The Biobank Access Decisions for FinnGen samples and data utilized in FinnGen Data Freeze 4 include: Auria Biobank AB17-5154, THL Biobank BB2017_55, BB2017_111, BB2018_19, BB_2018_34, Finnish Red Cross Blood Service Biobank 7.12.2017, Helsinki Biobank HUS/359/2017 and Northern Finland Biobank Borealis BB_2017_1013. The Ethical Review Board of the Hospital District of Helsinki and Uusimaa approved the FinnGen study protocol Nr HUS/990/2017. The FinnGen preparatory project is approved by THL, approval numbers THL/2031/6.02.00/2017, amendments THL/341/6.02.00/2018, THL/2222/6.02.00/2018, THL/1101/5.05.00/2017, VRK43431/2017-3, KELA 131/522/2018, and Statistics Finland TK-53-1041-17, and THL/283/6.02.00/2019. All DNA samples and data in this study were pseudonymized.

**FinnGen Data availability**

The FinnGen data may be accessed through Finnish Biobanks' FinnBB portal (www.finbb.fi) and THL Biobank data through THL Biobank (https://thl.fi/en/web/thl-biobank).

**FinnGen Code availability**

The full genotyping and imputation protocol for FinnGen is described at dx.doi.org/10.17504/protocols.io.nmndc5e

#### FIGURES

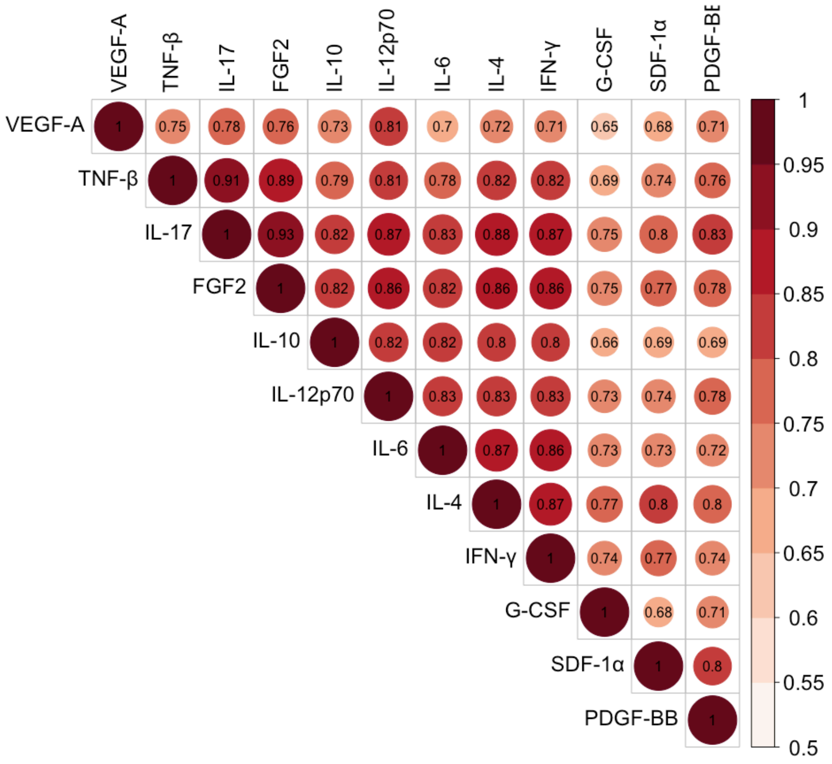


**Supplementary Figure 1. Correlation structure of the 12 inflammatory biomarkers.** The color and size of the circle represents the correlation of the inverse-rank normalized age and sex adjusted biomarkers. The Pearson correlations between the biomarkers ranged from 0.64 to 0.93, with a mean of 0.80.

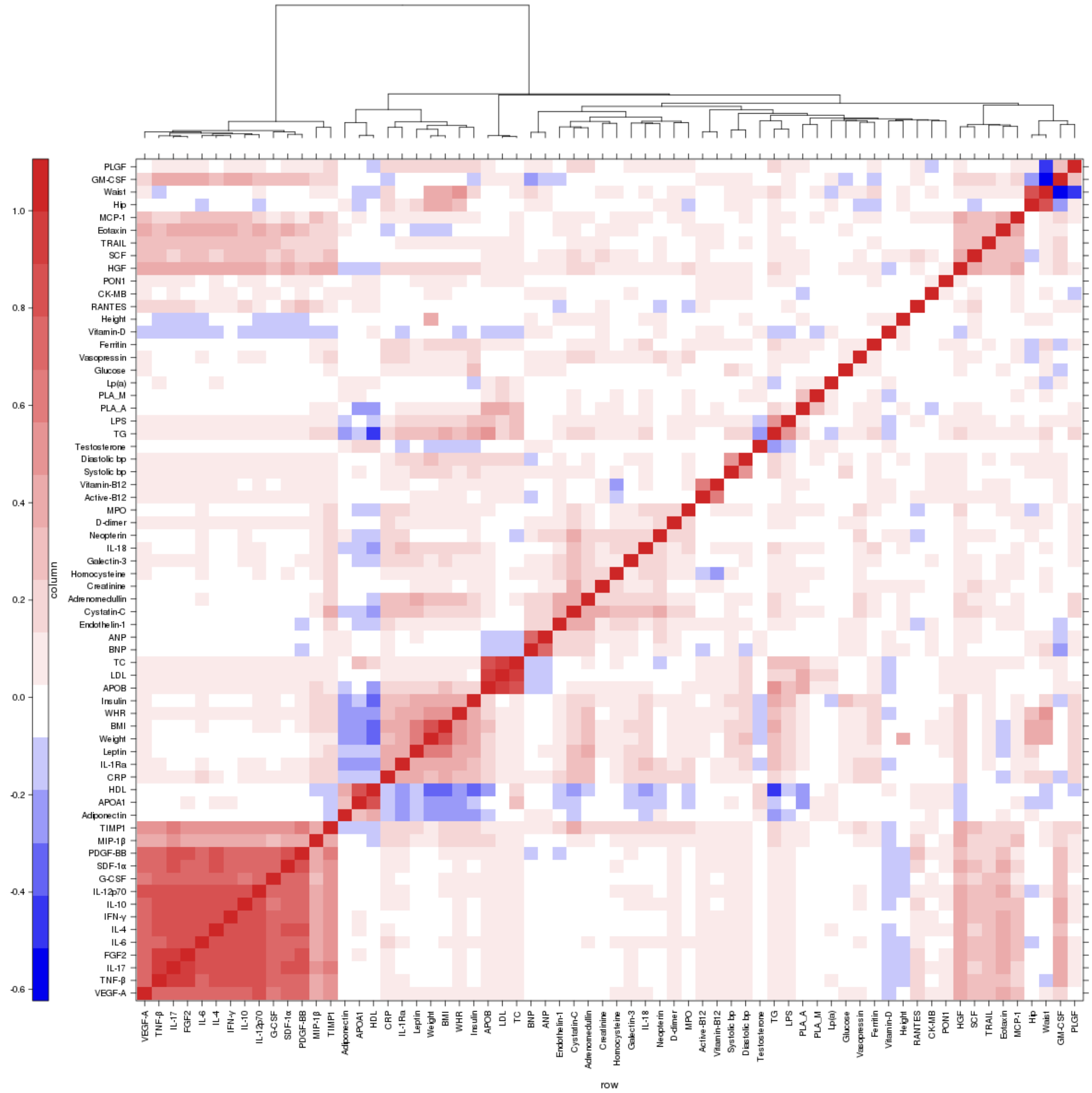


**Supplementary Figure 2. Correlation structure of the 66 quantitative traits.**

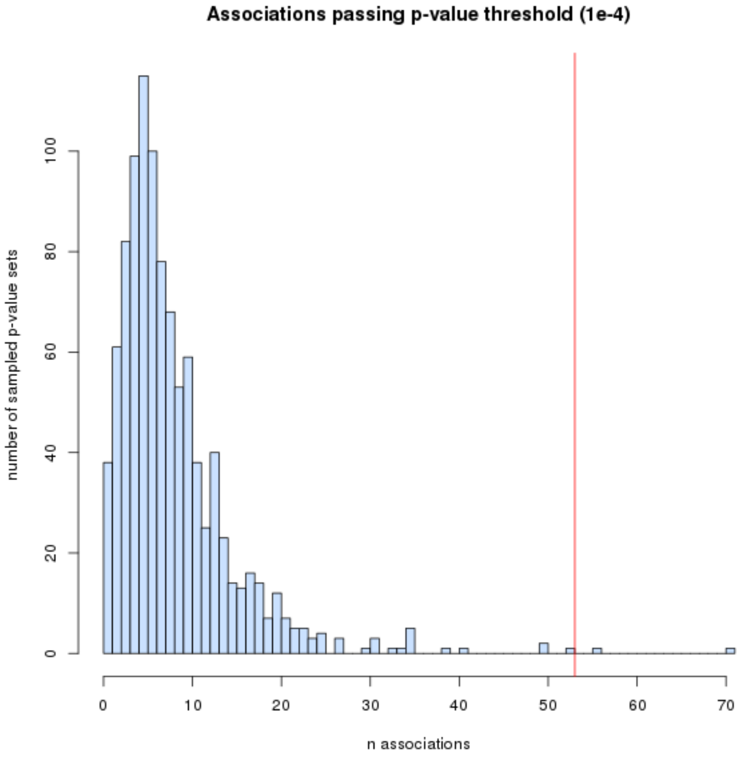


**Supplementary Figure 3. Empirical testing of chosen p-value threshold.** One thousand sets of 19 allele frequency-matched variants were sampled from 8.2 million non-coding variants. Allele frequencies were matched to the 19 putative casual variants selected to represent the 19 credible sets (Table 2). The number of associations passing the p-value threshold $1\times10^{-4}$ for each of these 1,000 sets of variants is plotted above. The median of this "null distribution" was 7 and 95% of the thousand sets had 20 associations or less passing the p-value threshold. The red line represents the number of significant associations observed for the 19 putative causal variants (n = 53). Only 0.3% of the thousand sampled sets had as many significant associations, indicating that the chosen p-value threshold is stringent.

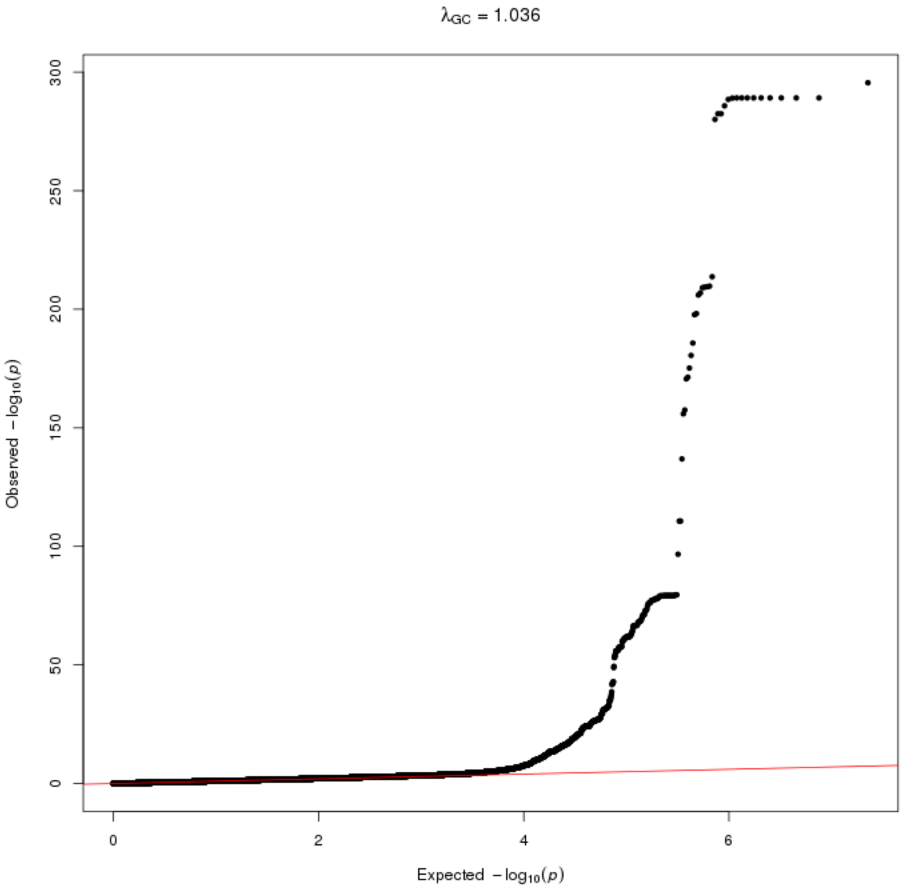


**Supplementary Figure 4. Quantile-quantile plot for multivariate GWAS results.**

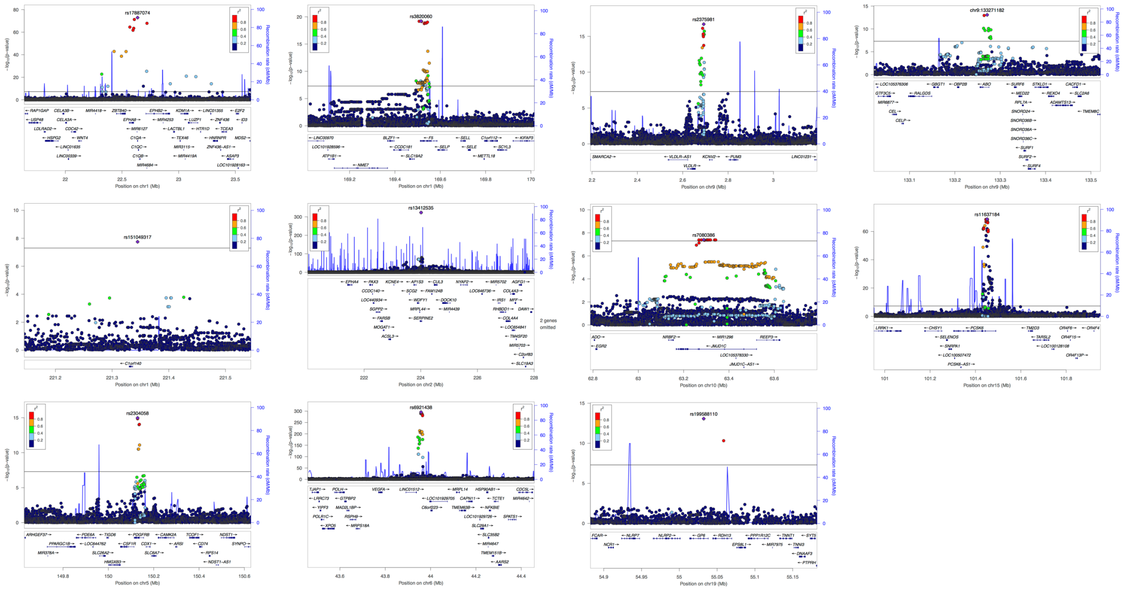

**Supplementary Figure 5. Locuszoom plots for each of the 11 genome-wide significant loci in the multivariate GWAS.**

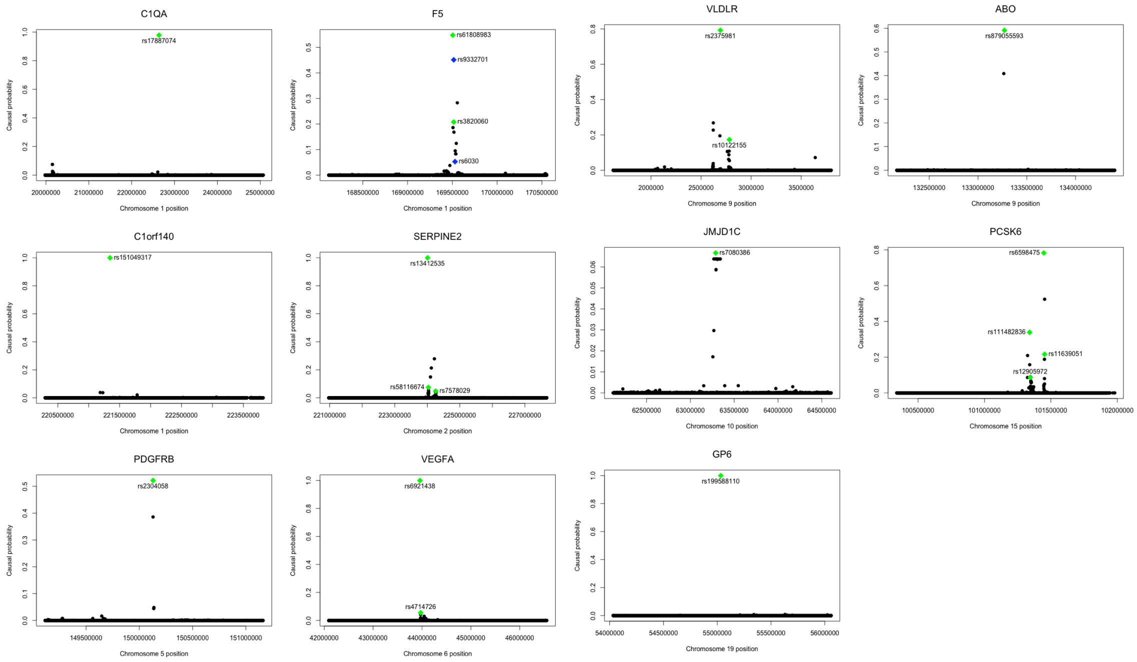


**Supplementary Figure 6**. **FINEMAP results of the 11 loci.** The SNP-wise causal probability is plotted on the y-axis and the chromosomal position on the x-axis. The initial representative variants are represented as green diamonds with accompanying rsids, and missense variants in high LD with them are represented as blue diamonds.

### TABLES

| Trait | Description |
|---|---|
| Systolic bp | Systolic blood pressure |
| Diastolic bp | Diastolic blood pressure |
| HDL | High density lipoprotein |
| TG | Triglycerides |
| TC | Total cholesterol |
| LDL | Low density lipoprotein |
| Lp(a) | Lipoprotein (a) |
| APOA1 | Apolipoprotein A-I |
| APOB | Apolipoprotein B |
| Galectin-3 | Galectin-3 |
| LPS | Lipopolysaccharide |
| CRP | C-reactive protein |
| HGF | Hepatocyte growth factor |
| SCF | Stem cell factor |
| SDF-1α | Stromal cell derived factor 1 alpha (CXCL12) |
| TNF-β | Tumor necrosis factor beta |
| TRAIL | TNF related apoptosis inducing ligand |
| IL-4 | Interleukin-4 |
| IL-6 | Interleukin-6 |
| IL-10 | Interleukin-10 |
| IL-12p70 | Interleukin-12p70 |
| IL-17 | Interleukin-17 |
| Eotaxin | Eotaxin (CCLL11) |
| FGF2 | Basic fibroblast growth factor |
| G-CSF | Granulocyte colony stimulating factor |
| GM-CSF | Granulocyte monocyte colony stimulating factor |
| IFN-γ | Interferon gamma |
| MCP-1 | Monocyte chemoattractant protein 1 (CCL2) |
| PDGF-BB | Platelet derived growth factor BB |
| MIP-1β | Macrophage inflammatory protein 1 beta (CCL4) |
| RANTES | Regulated on Activation, Normal T cell Expressed and Secreted (CCL5) |
| VEGF-A | Vascular endothelial cell growth factor A |
| Active-B12 | Active vitamin B12 |
| Adiponectin | Adiponectin |
| BNP | Brain natriuretic peptide |
| CK-MB | Creatine kinase isoenzyme MB |
| Creatinine | Creatinine |
| Vasopressin | C-terminal pro-vasopressin |
| Endothelin-1 | C-terminal pro-endothelin 1 |
| Cystatin-C | Cystatin C |
| D-dimer | D-dimer |
| Ferritin | Ferritin |
| Homocysteine | Homocysteine |
| IL-18 | Interleukin-18 |
| IL-1Ra | Interleukin-1 receptor antagonist |
| Leptin | Leptin |
| MPO | Myeloperoxidase |
| Adrenomedullin | Mid-regional pro-adrenomedullin |
| ANP | Mid-regional pro-atrial natriuretic peptide |
| Neopterin | Neopterin |
| PLA_M | Phospholipase A2 mass |
| PLA_A | Phospholipase A2 activity |
| PLGF | Placental growth factor |
| PON1 | Paraoxonase 1 |
| TIMP1 | Tissue inhibitor metalloproteinase 1 |
| Glucose | Glucose corrected for fasting |
| Insulin | Insulin corrected for fasting |
| Testosterone | Testosterone |
| Vitamin-B12 | Vitamin B12 (Cobalamin) |
| Vitamin-D | Vitamin D |
| BMI | Body Mass Index |
| Waist | Waist circumference |
| Hip | Hip circumference |
| WHR | Waist hip ratio |
| Weight | Weight |
| Height | Height |

**Supplementary Table 1: List of the 66 quantitative traits.**

| Variant[a] | Locus | AF[b] (FIN enrichment) | Most severe consequence | LD with lead variant ($r^2$) | Multivariate p-value | Minimum univariate p-value (biomarker) | Driver traits | Novel biomarker association[c] | FinnGen disease associations[d] | FinnGen association statistics | | Novel disease association[e] |
|---|---|---|---|---|---|---|---|---|---|---|---|---|
| | | | | | | | | | | OR | p-value | |
| rs45498698 | *C1QA* | 1.61% (1.70) | missense variant | 0,87 | 2,97E-65 | 6.39E-23 (TNF-β) | TNF-β | NO | — | — | — | — |
| rs144329757 | *C1QA* | 1.56% (1.64) | missense variant | 0,88 | 1,97E-62 | 2.54E-20 (TNF-β) | TNF-β | NO | — | — | — | — |
| **rs17887074** | *C1QA* | 1.48% (4.64) | missense variant | 1 | 1,21E-73 | 1.70E-23 (TNF-β) | TNF-β | NO | — | — | — | — |
| rs6027 | *F5* | 8,26 % | missense variant | 0,21 | 6,06E-09 | 8.83E-5 (IL-4) | IL-4 | YES | Venous thromboembolism<br>DVT of lower extremities<br>DVT of lower extremities and pulmonary embolism | 1.17<br>1.27<br>1.18 | 1.8E-6<br>7.8E-8<br>1.5E-6 | NO |
| | | | | | | | | | Diseases of veins, lymphatic vessels and lymph nodes, not elsewhere classified | 1,11 | 2.4E-5 | NO |
| rs6030 | *F5* | 29,71 % | missense variant | 0,997 | 1,78E-19 | 1.39E-3 (VEGF-A) | IL-4, !L-12 | YES | — | — | — | — |
| rs6032* | *F5* | 21,97 % | missense variant | 0,63 | 4,60E-09 | 3.14E-4 (VEGF-A) | IL-4 | YES | Venous thromboembolism<br>Phlebitis and thrombophlebitis<br>DVT of lower extremities and pulmonary embolism | 0.86<br>0.87<br>0.86 | 1.3E-11<br>6.9E-5<br>2.8E-11 | NO |
| | | | | | | | | | Hypo-osmolality and hyponatraemia | 1,18 | 9.5E-5 | YES |
| rs12694627 | *SERPINE2* | 74,75 % | splice region variant | 0,083 | 4,55E-13 | 5.31E-4 (IL-10) | PDGF-BB, SDF-1α | NO | — | — | — | — |
| rs3738952 | *SERPINE2* | 9,78 % | missense variant | 0,013 | 1,21E-08 | 6.27E-3 (IL-10) | PDGF-BB | NO | — | — | — | — |
| rs115402675 | *VEGFA* | 6.49% (2.12) | missense variant | 0,022 | 1,95E-12 | 1.65E-4 (VEGF-A) | VEGF-A | NO | — | — | — | — |
| rs57288791 | *VEGFA* | 8.83% (1.60) | frameshift variant | 0,023 | 5,02E-11 | 6.49E-6 (VEGF-A) | VEGF-A | NO | — | — | — | — |
| **rs199588110** | *GP6* | 0.33% (3.69) | missense variant | 1 | 8,54E-14 | 1.25E-17 (IL-17) | all 12 biomarkers | NO | Benign neoplasm of meninges | 6,4 | 4.9E-5 | YES |

**Supplementary Table 2: Results of the 13 functional variants reaching genome-wide significance in the multivariate GWAS.**

* represents two other missense variants (rs4525 and rs4524) in high LD ($r^2 > 0.98$) with the same disease associations, excluding the association with hypo-osmolality and hyponatraemia that was only observed for rs6032.

[a] Bolded variants are lead variants.

[b] AF = allele frequency, FIN enrichment = AF in Finns compared to AF in Non-Finnish Europeans excluding Estonians in the gnomAD genomes database; reported if it was at least 1.5-fold.

[c] Previous associations with the 12 biomarkers were searched for in the NHGRI-EBI GWAS Catalog within a region encompassing ±500 kB around the variant. An association was regarded novel if no associations with any of the 12 biomarkers had been reported in this region.

[d] Only associations that remain significant after conditioning are reported here. Closely related disease diagnoses are represented in a shared cell and their replication is assessed jointly. DVT = deep vein thrombosis.

[e] Novelty of disease associations was assessed at gene-level.

LD = linkage disequilibrium

**1 credible set**

| Cred1 | Prob1 |
|---|---|
| *C1QA* | |
| chr1:22637683:G:A | 0,975 |
| C1orf140 | |
| chr1:221344914:A:G | 1,000 |
| *ABO* | |
| chr9:133271182:T:C | 0,578 |
| chr9:133264504:G:GAAACTGCC | 0,422 |
| *PDGFRB* | |
| chr5:150128981:C:G | 0,546 |
| chr5:150128912:C:T | 0,405 |
| *JMJD1C* | |
| chr10:63288546:C:A | 0,064 |
| chr10:63306426:G:T | 0,062 |
| chr10:63304084:T:A | 0,062 |
| chr10:63302248:G:C | 0,0621003 |
| chr10:63318234:C:G | 0,062 |
| chr10:63304076:ACTTTGCC:A | 0,062 |
| chr10:63341447:G:C | 0,062 |
| chr10:63336490:C:T | 0,062 |
| chr10:63303060:C:T | 0,062 |
| chr10:63267383:G:T | 0,062 |
| chr10:63290899:A:G | 0,062 |
| chr10:63278270:G:C | 0,062 |
| chr10:63311455:C:A | 0,062 |
| chr10:63292782:C:T | 0,058 |
| chr10:63292445:T:C | 0,058 |
| chr10:63267850:A:T | 0,042 |
| *GP6* | |
| chr19:55032292:G:A | 1,000 |

**2 credible sets**

| Cred1 | Prob1 | Cred2 | Prob2 |
|---|---|---|---|
| *F5* | | | |
| chr1:169515314:T:G | 0,175 | chr1:169505159:C:T | 0,533 |
| chr1:169543742:G:A | 0,164 | chr1:169515529:A:G | 0,461 |
| chr1:169507088:G:A | 0,148 | | |
| chr1:169517529:T:G | 0,141321 | | |
| chr1:169531818:T:C | 0,117 | | |
| chr1:169538606:G:T | 0,103 | | |
| chr1:169537003:A:G | 0,103489 | | |
| *VEGFA* | | | |
| chr6:43957870:G:A | 1,000 | chr6:43974045:A:G | 0,055 |
| | | chr6:43974829:T:C | 0,053 |
| | | chr6:43973361:A:T | 0,053 |
| | | chr6:43973945:T:C | 0,053 |
| | | chr6:43972634:A:AG | 0,053 |
| | | chr6:43974508:A:G | 0,053 |
| | | chr6:43974249:C:T | 0,053 |
| | | chr6:43974275:A:T | 0,053 |
| | | chr6:43973612:A:G | 0,053 |
| | | chr6:43972532:A:G | 0,053 |
| | | chr6:43972465:A:G | 0,053 |
| | | chr6:43972186:G:T | 0,053 |
| | | chr6:43972253:A:G | 0,053 |
| | | chr6:43973908:C:T | 0,053 |
| | | chr6:43974487:T:C | 0,052 |
| | | chr6:43972413:C:T | 0,050 |
| | | chr6:43974652:T:C | 0,042 |
| | | chr6:43971194:A:G | 0,041 |
| | | chr6:43971636:A:G | 0,018 |
| *VLDLR* | | | |
| chr9:2692583:C:G | 0,748 | chr9:2782458:A:G | 0,162 |
| chr9:2687795:A:T | 0,234 | chr9:2780496:C:CT | 0,109 |
| | | chr9:2763894:C:A | 0,109 |
| | | chr9:2762262:T:A | 0,109 |
| | | chr9:2762263:C:A | 0,109 |
| | | chr9:2777384:T:A | 0,092 |
| | | chr9:2774758:A:G | 0,067 |
| | | chr9:2782774:T:C | 0,061 |
| | | chr9:2776401:C:G | 0,023 |
| | | chr9:2787594:C:G | 0,020 |
| | | chr9:2777062:G:T | 0,017 |
| | | chr9:2787856:A:C | 0,017 |
| | | chr9:2791828:T:C | 0,011 |
| | | chr9:2791753:A:T | 0,010 |
| | | chr9:2792552:A:C | 0,009 |
| | | chr9:2793342:A:T | 0,006 |
| | | chr9:2797082:G:A | 0,006 |
| | | chr9:2796867:G:A | 0,006 |
| | | chr9:2794338:T:C | 0,005 |
| | | chr9:2795985:G:C | 0,004 |
| | | chr9:2796135:CCA:C | 0,004 |

**3 credible sets**

| Cred1 | Prob1 | Cred2 | Prob2 | Cred3 | Prob3 |
|---|---|---|---|---|---|
| *SERPiNE2* | | | | | |
| chr2:224010157:G:A | 0,998104 | chr2:224035992:CA:C | 0,067 | chr2:224257750:T:A | 0,046 |
| | | chr2:224038180:A:G | 0,050 | chr2:224260063:A:G | 0,045 |
| | | chr2:224041598:C:T | 0,040 | chr2:224261196:C:T | 0,043 |
| | | chr2:224034475:A:G | 0,037 | chr2:224263607:A:G | 0,041 |
| | | chr2:224036383:A:G | 0,036 | chr2:224261500:A:T | 0,041 |
| | | chr2:224036374:A:G | 0,036 | chr2:224263458:T:G | 0,040 |
| | | chr2:224028228:A:G | 0,036 | chr2:224258053:G:C | 0,040 |
| | | chr2:224036011:C:A | 0,033 | chr2:224259370:T:C | 0,040 |
| | | chr2:224035127:T:C | 0,032 | chr2:224252531:C:T | 0,040 |
| | | chr2:224033438:C:T | 0,032 | chr2:224251564:C:G | 0,040 |
| | | chr2:224035435:A:T | 0,032 | chr2:224251491:A:G | 0,040 |
| | | chr2:224035116:T:TG | 0,032 | chr2:224255099:A:G | 0,039 |
| | | chr2:224035358:T:C | 0,032 | chr2:224268165:G:A | 0,036 |
| | | chr2:224035405:CAG:C | 0,032 | chr2:224263185:C:T | 0,034 |
| | | chr2:224033029:C:T | 0,032 | chr2:224268510:G:C | 0,032 |
| | | chr2:224035198:G:A | 0,032 | chr2:224229124:C:T | 0,018 |
| | | chr2:224035419:T:G | 0,032 | chr2:224230121:C:T | 0,012 |
| | | chr2:224031620:C:T | 0,032 | chr2:224229721:C:CA | 0,012 |
| | | chr2:224031625:C:A | 0,032 | chr2:224229749:G:A | 0,012 |
| | | chr2:224035656:A:T | 0,030 | chr2:224236034:A:G | 0,011 |
| | | chr2:224035622:G:T | 0,030 | chr2:224228932:C:T | 0,011 |
| | | chr2:224026230:G:A | 0,030 | chr2:224229526:A:T | 0,011 |
| | | chr2:224038091:A:T | 0,029 | chr2:224230255:G:A | 0,011 |
| | | chr2:224036103:C:T | 0,029 | chr2:224229279:G:A | 0,011 |
| | | chr2:224032508:C:T | 0,026 | chr2:224250193:GAAACCCCATCACTACTAA:G | 0,010 |
| | | chr2:224035885:T:C | 0,026 | chr2:224239981:C:T | 0,010 |
| | | chr2:224029381:TA:T | 0,026 | chr2:224237902:G:C | 0,010 |
| | | chr2:224033710:A:G | 0,025 | chr2:224242921:A:G | 0,010 |
| | | chr2:224031721:A:G | 0,025 | chr2:224234466:G:T | 0,010 |
| | | | | chr2:224243642:G:A | 0,010 |
| | | | | chr2:224239836:G:C | 0,010 |
| | | | | chr2:224239156:G:T | 0,010 |
| | | | | chr2:224241641:T:C | 0,010 |
| | | | | chr2:224247045:C:T | 0,010 |
| | | | | chr2:224240716:A:G | 0,010 |
| | | | | chr2:224241834:C:T | 0,010 |
| | | | | chr2:224234172:C:G | 0,010 |
| | | | | chr2:224205442:C:T | 0,009 |
| | | | | chr2:224226428:C:T | 0,008 |
| | | | | chr2:224227446:A:G | 0,008 |
| | | | | chr2:224220047:G:A | 0,008 |
| | | | | chr2:224216258:T:G | 0,008 |
| | | | | chr2:224216965:C:T | 0,008 |
| | | | | chr2:224209822:A:G | 0,008 |
| | | | | chr2:224202443:G:C | 0,008 |
| | | | | chr2:224181156:C:T | 0,008 |
| | | | | chr2:224175988:AT:A | 0,008 |
| | | | | chr2:224198340:C:T | 0,006 |
| | | | | chr2:224272987:A:G | 0,006 |
| | | | | chr2:224229118:T:C | 0,005 |
| | | | | chr2:224270805:GTTAAC:G | 0,005 |
| | | | | chr2:224275991:AG:A | 0,004 |
| | | | | chr2:224275258:A:G | 0,004 |
| | | | | chr2:224280597:C:T | 0,004 |

**4 credible sets**

| Cred1 | Prob1 | Cred2 | Prob2 | Cred3 | Prob3 | Cred4 | Prob4 |
|---|---|---|---|---|---|---|---|
| *PCSK6* | | | | | | | |
| chr15:101452793:C:A | 0,321 | chr15:101446425:G:T | 0,704 | chr15:101345122:G:A | 0,127 | chr15:101339772:G:A | 0,537 |
| chr15:101451543:G:C | 0,296 | chr15:101451568:T:C | 0,082 | chr15:101345382:A:G | 0,122 | chr15:101340399:T:A | 0,325 |
| chr15:101452832:A:G | 0,252 | chr15:101446466:C:T | 0,065 | chr15:101348606:T:C | 0,109 | chr15:101335494:G:T | 0,058 |
| chr15:101451951:A:G | 0,100 | chr15:101446770:G:A | 0,058 | chr15:101352810:G:A | 0,094 | chr15:101333461:T:C | 0,058 |
| | | chr15:101447186:T:C | 0,032 | chr15:101351713:GT:G | 0,094 | | |
| | | chr15:101451106:A:G | 0,020 | chr15:101351332:T:A | 0,093 | | |
| | | | | chr15:101351296:A:G | 0,093 | | |
| | | | | chr15:101350243:C:A | 0,093 | | |
| | | | | chr15:101350358:T:A | 0,093 | | |
| | | | | chr15:101346648:A:G | 0,0689712 | | |

**Supplementary Table 3: Variants within the 19 fine-mapped credible sets.** Missense variants are highlighted with blue font.

| N | *C1QA* | *F5* | *C1orf140* | *SERPINE2* | *PDGFRB* | *JMJD1C* | *VEGFA* | *VLDLR* | *ABO* | *PCSK6* | *GP6* |
|---|---|---|---|---|---|---|---|---|---|---|---|
| 1 | 0.451 | 0 | **0.541** | 0 | **0.591** | **0.796** | 0 | 0.00148 | **0.652** | 0 | **0.688** |
| 2 | **0.473** | 0.251 | 0.42 | 0.00164 | 0.399 | 0.206 | **0.575** | 0.237 | 0.348 | 0.00253 | 0.303 |
| 3 | 0.0766 | **0.602** | 0.0388 | 0.324 | 0.01 | 0 | 0.424 | **0.586** | 0 | 0.21 | 0 |
| 4 | 0 | 0.147 | 0 | **0.671** | 0 | 0 | 0.00137 | 0.173 | 0 | **0.441** | 0 |
| 5 | 0 | 0 | 0 | 0.00311 | 0 | 0 | 0 | 0.00293 | 0 | 0.322 | 0 |
| 6 | 0 | 0 | 0 | 0 | 0 | 0 | 0 | 0 | 0 | 0.024 | 0 |
| 7-10 | 0 | 0 | 0 | 0 | 0 | 0 | 0 | 0 | 0 | 0 | 0 |

**Supplementary Table 4: Posterior probabilities estimated by multivariate FINEMAP for the number of causal signals within each locus**

| *Locus* | # Conditioning rounds | # Credible sets from FINEMAP | # FINEMAP variants in conditional analysis (%) | Conditioned variants |
|---|---|---|---|---|
| *C1QA* | 1 | 1 | 1 (100) | chr1:22637683:G:A |
| *F5* | 2 | 2 | 2 (100) | chr1:169515314:T:G, chr1:169505159:C:T:C |
| *C1orf140* | 1 | 1 | 1 (100) | chr1:221344914:A:G |
| *SERPINE2* | 3 | 3 | 1 (33,3) | chr2:224010157:G:A, chr2:224041598:C:T, chr2:224261196:C:T |
| *PDGFRB* | 1 | 1 | 1 (100) | chr5:150128981:C:G |
| *VEGFA* | 2 | 2 | 2 (100) | chr6:43957870:G:A, chr6:43974045:A:G:A |
| *VLDLR* | 1 | 2 | 1 (50) | chr9:2692583:C:G |
| *ABO* | 1 | 1 | 1 (100) | chr9:133271182:T:C |
| *JMJD1C* | 1 | 1 | 1 (100) | chr10:63288546:C:A:C |
| *PCSK6* | 2 | 4 | 1 (25) | chr15:101451543:G:C:G, chr15:101446425:G:T:G |
| *GP6* | 1 | 1 | 1 (100) | chr19:55032292:G:A:G |
| ***Total*** | **16** | **19** | **13 (68,4)** | |

**Supplementary Table 5: Comparison of FINEMAP and conditional analysis results.**

| Variant[a] | Locus | AF[b] (FIN enrichment) | FinnGen disease associations[c] | FinnGen association statistics | | | Replication in UKBB[d] | UKBB association statistics | | | Novel disease association[e] |
|---|---|---|---|---|---|---|---|---|---|---|---|
| | | | | OR | p-value | № cases | | OR | p-value | № cases | |
| rs6027* | *F5* | 8.26% | Venous thromboembolism<br>DVT of lower extremities<br>DVT of lower extremities and pulmonary embolism | 1.17<br>1.27<br>1.18 | 1.8E-6<br>7.8E-8<br>1.5E-6 | 6,913<br>3,592<br>6,019 | Phlebitis and thrombophlebitis | 1.06 | 0.27 | 3,900 | NO |
| | | | Diseases of veins, lymphatic vessels and lymph nodes, not elsewhere classified | 1.11 | 2.4E-5 | 19,930 | No replication phenotype | — | — | — | NO |
| rs6032** | *F5* | 21.97% | Venous thromboembolism<br>Phlebitis and thrombophlebitis<br>DVT of lower extremities and pulmonary embolism | 0.86<br>0.87<br>0.86 | 1.3E-11<br>6.9E-5<br>2.8E-11 | 6,913<br>2,506<br>6,019 | Phlebitis and thrombophlebitis | 0.82 | 3.4E-14 | 3,900 | NO |
| | | | Hypo-osmolality and hyponatraemia | 1.18 | 9.5E-5 | 1,648 | Hypo-osmolality and hyponatraemia | 1.01 | 0.78 | 1,826 | YES |
| **rs13412535** | *SERPINE2* | 19.8% | Hypertrophic scar | 1.34 | 7.5E-5 | 591 | Keloid scar<br>Aquired keratoderma | 0.90<br>1.52 | 0.35<br>0.02 | 216<br>87 | YES |
| **rs550057** | *ABO* | 31.0% | Anemias<br>Other and unspecified anaemias<br>Other anaemias<br>Diseases of the blood and blood-forming organs | 1.12<br>1.10<br>1.11<br>1.06 | 4.7E-5<br>4.9E-5<br>2.6E-5<br>2.9E-5 | 9,145<br>4,279<br>4,337<br>14,375 | Other anemias<br>Red boold cell count | 0.99<br>NA | 0.71<br>1.26E-212 | 12,256<br>445,000 | NO |
| | | | Visual field defects<br>Diseases of the eye and adnexa | 1.24<br>1.04 | 4.4E-5<br>9.4E-6 | 885<br>58,498 | Visual field defects | 1.00 | 0.97 | 334 | YES |
| | | | Diseases of the ear and mastoid process | 1.04 | 4.8E-5 | 31,579 | No replication phenotype | — | — | — | YES |
| **rs199588110*** | *GP6* | 0.33% (3.69) | Benign neoplasm of meninges | 6.40 | 4.9E-5 | 934 | Benign neoplasm of brain, cranial nerves, meninges | 0.09 | 0.31 | 774 | YES |

**Supplementary Table 6: Replication of FinnGen disease associations in the UKBB.**

* missense variant

**represents two other missense variants (rs4525 and rs4524) in high LD ($r^2 > 0.98$) with the same disease associations, excluding the association with hypo-osmolality and hyponatraemia that was only observed for rs6032.

[a] Bolded variants are putative causal variants identified by fine-mapping.

[b] AF = allele frequency, FIN enrichment = AF in Finns compared to AF in Non-Finnish Europeans excluding Estonians in the gnomAD genomes database; reported if it was at least 1.5-fold.

[c] Only associations that remain significant after conditioning are reported here. Closely related disease diagnoses are represented in a shared cell and their replication is assessed jointly. DVT = deep vein thrombosis.

[d] Replication in the UKBB was analyzed using the summary statistics of SAIGE analysis on ~400,000 samples run by the Lee lab, University of Michigan, https://www.leelabsg.org/resources and the study by Kichaev[2] .

[e] Novelty of disease associations was assessed at gene-level.

| Variant | Locus | QTL type | Gene (tissue) | p-value | Study | high-LD QTL lead variant ($r^2$) |
|---|---|---|---|---|---|---|
| rs17887074 | *C1QA* | | | | | |
| | | – | – | – | – | – |
| rs3820060 | *F5* | | | | | |
| | | eQTL | F5 (whole blood) | 9,2E-118 | GTEX7 | |
| | | eQTL | NME7 (whole blood) | 1,8E-10 | GTEX7 | |
| | | high-LD pQTL | CAMK1 | 2,0E-110 | Suhre | rs4525 (0.70) |
| | | high-LD pQTL | SEC13 | 2,3E-12 | Suhre | rs1800594 (0.96) |
| | | high-LD pQTL | NPTX2 | 3,4E-14 | Suhre | rs9287090 (0.70) |
| | | high-LD pQTL | SIG11 | 9,3E-97 | Suhre | rs9287090 (0.70) |
| | | high-LD pQTL | TFPI | 3,2E-24 | Suhre | rs10800453 (0.70) |
| rs9332701 | *F5* | | | | | |
| | | pQTL | F5 | 1,0E-23 | Suhre | |
| | | high-LD eQTL | NME7 (blood) | 1,4E-27 | GTEX7 | rs61808983 (0.97) |
| | | high-LD pQTL | EHBP1 | 7,1E-18 | Sun | rs61808983 (0.97) |
| rs151049317 | *C1orf140* | | | | | |
| | | – | – | – | – | |
| rs13412535 | *SERPINE2* | | | | | |
| | | pQTL | PDGF-BB | 4,6E-13 | Sun | |
| | | high-LD pQTL | SERPINE2 | 1,1E-46 | Sun | rs68066031 (0.99) |
| | | high-LD pQTL | SERPINE2 | 5,9E-246 | Emilsson | rs68066031 (0.99) |
| rs58116674 | *SERPINE2* | | | | | |
| | | – | – | – | – | |
| rs7578029 | *SERPINE2* | | | | | |
| | | – | – | – | – | |
| rs2304058 | *PDGFRB* | | | | | |
| | | pQTL | PDGFRB | 2.3E-458 | Sun | |
| | | high-LD pQTL | PDGFRB | 1,0E-300 | Emilsson | rs4705415 (0.70) |
| | | high-LD pQTL | PDGFRB | 5,0E-161 | Suhre | rs2240781 (0.95) |
| | | high-LD pQTL | PDGFRB | 1,0E-33 | Sasayama | rs740750 (0.99) |
| rs6921438 | *C6orf223 / VEGFA* | | | | | |
| | | pQTL | VEGFA | 7,8E-71 | Sun | |
| | | pQTL | VEGFA isoform 121 | 1,7E-234 | Sun | |
| rs4714726 | *C6orf223 / VEGFA* | | | | | |
| | | – | – | – | – | – |
| rs2375981 | *VLDLR / KCNV2* | | | | | |
| | | – | – | – | – | – |
| rs10122155 | *VLDLR / KCNV2* | | | | | |
| | | – | – | – | – | – |
| rs550057 | *ABO* | | | | | |
| | | pQTL | ALPI | 2,8E-19 | Sun | |
| | | pQTL | CHST15 | 1,0E-30 | Sun | |
| | | pQTL | FAM177A1 | 9,3E-19 | Sun | |
| | | pQTL | JAG1 | 8,3E-14 | Sun | |
| rs7080386 | *JMJD1C* | | | | | |
| | | pQTL | HB-EGF | 1,6E-13 | Sun | |
| | | high-LD eQTL | NRBF2 (thyroid) | 2,5E-07 | GTEX7 | rs4595427 (0.81) |
| | | high-LD eQTL | NRBF2 (whole blood) | 4,6E-06 | GTEX7 | rs10995477 (0.80) |
| | | high-LD eQTL | NRBF2 (brain frontal cortex BA9) | 2,2E-09 | GTEX7 | rs10761729 (0.79) |
| rs111482836 | *PCSK6* | | | | | |
| | | – | – | – | – | – |
| rs12905972 | *PCSK6* | | | | | |
| | | – | – | – | – | – |
| rs6598475 | *PCSK6* | | | | | |
| | | – | – | – | – | – |
| rs11639051 | *PCSK6* | | | | | |
| | | – | – | – | – | – |
| rs199588110 | *GP6* | | | | | |
| | | – | – | – | – | – |

**Supplementary Table 7: Expression and protein quantitative trait loci of the 19 putative causal variants.**

| Variant (rsid) | Locus | Putative causal/Functional | Disease | N (cases/controls) | OR (P-value) | | LD with FG lead |
|---|---|---|---|---|---|---|---|
| | | | | | Original | Conditinal | |
| chr1:169514323:T:C (rs6027) | *F5* | Functional | Diseases of veins, lymphatic vessels and lymph nodes, not elsewhere classified (FINNGEN) | 19930/97214 | 1,11 (2,42E-05) | 1,12 (2,19E-06) | 0,002 |
| | | | Diseases of veins, lymphatic vessels and lymph nodes, not elsewhere classified | 22948/153951 | 1,09 (3,64E-05) | 1,10 (3,86E-06) | 0,002 |
| | | | DVT of lower extremities and pulmonary embolism | 6019/170880 | 1,18 (1,47E-06) | 1,22 (1,22E-08) | 0,002 |
| | | | DVT of lower extremities | 3592/153951 | 1,27 (7,80E-08) | 1,32 (5,65E-10) | 0,002 |
| | | | Venous thromboembolism | 6913/169986 | 1,17 (1,81E-06) | 1,21 (8,41E-09) | 0,002 |
| chr1:169542317:T:C (rs6032) | *F5* | Functional | Hypo-osmolality and hyponatraemia | 1648/160461 | 1,18 (9,49E-05) | 0,94 (7,86E-01) | 0,965 |
| | | | Pulmonary heart disease | 3016/173883 | 0,85 (8,03E-07) | 0,87 (7,13E-06) | 0,008 |
| | | | DVT of lower extremities and pulmonary embolism | 6019/170880 | 0,86 (2,78E-11) | 0,87 (6,00E-09) | 0,008 |
| | | | Phlebitis and thrombophlebitis (not including DVT) | 2506/153951 | 0,87 (6,92E-05) | 0,90 (2,29E-03) | 0,008 |
| | | | DVT of lower extremities | 3592/153951 | 0,86 (2,32E-07) | 0,90 (3,35E-04) | 0,008 |
| | | | Pulmonary embolism | 3016/173597 | 0,85 (8,43E-07) | 0,87 (6,92E-06) | 0,008 |
| | | | Pulmonary heart disease, diseases of pulmonary circulation | 3302/173597 | 0,87 (7,89E-06) | 0,89 (7,23E-05) | 0,008 |
| | | | Venous thromboembolism | 6913/169986 | 0,86 (1,25E-11) | 0,88 (4,16E-09) | 0,008 |
| | | | Other ILD-related CVD-co-morbidities | 3260/114642 | 0,87 (1,67E-05) | 0,88 (3,35E-05) | 0,008 |
| chr1:169542496:T:C (rs4525) | *F5* | Functional | Pulmonary heart disease | 3016/173883 | 0,86 (1,10E-06) | 0,87 (9,68E-06) | 0,008 |
| | | | DVT of lower extremities and pulmonary embolism | 6019/170880 | 0,86 (2,50E-11) | 0,87 (5,47E-09) | 0,008 |
| | | | Phlebitis and thrombophlebitis (not including DVT) | 2506/153951 | 0,87 (4,92E-05) | 0,90 (1,76E-03) | 0,008 |
| | | | DVT of lower extremities | 3592/153951 | 0,86 (2,03E-07) | 0,90 (3,01E-04) | 0,008 |
| | | | Pulmonary embolism | 3016/173597 | 0,86 (1,16E-06) | 0,87 (9,40E-06) | 0,008 |
| | | | Pulmonary heart disease, diseases of pulmonary circulation | 3302/173597 | 0,87 (1,08E-05) | 0,89 (9,62E-05) | 0,008 |
| | | | Venous thromboembolism | 6913/169986 | 0,86 (1,01E-11) | 0,88 (3,43E-09) | 0,008 |
| | | | Other ILD-related CVD-co-morbidities | 3260/114642 | 0,88 (2,78E-05) | 0,88 (6,87E-05) | 0,008 |
| chr1:169542517:T:C (rs4524) | *F5* | Functional | Pulmonary heart disease | 3016/173883 | 0,85 (7,91E-07) | 0,87 (6,76E-06) | 0,008 |
| | | | DVT of lower extremities and pulmonary embolism | 6019/170880 | 0,86 (1,95E-11) | 0,87 (3,94E-09) | 0,008 |
| | | | Phlebitis and thrombophlebitis (not including DVT) | 2506/153951 | 0,87 (5,79E-05) | 0,90 (1,90E-03) | 0,008 |
| | | | DVT of lower extremities | 3592/153951 | 0,86 (1,84E-07) | 0,90 (2,81E-04) | 0,008 |
| | | | Pulmonary embolism | 3016/173597 | 0,85 (8,34E-07) | 0,86 (6,57E-06) | 0,008 |
| | | | Pulmonary heart disease, diseases of pulmonary circulation | 3302/173597 | 0,87 (8,83E-06) | 0,89 (7,77E-05) | 0,008 |
| | | | Venous thromboembolism | 6913/169986 | 0,86 (8,00E-12) | 0,88 (2,57E-09) | 0,008 |
| | | | Other ILD-related CVD-co-morbidities | 3260/114642 | 0,87 (1,91E-05) | 0,88 (3,58E-05) | 0,008 |
| chr2:224010157:G:A (rs13412535) | *SERPINE2* | Putative causal | Hypertrophic scar | 591/168348 | 1,34 (7,51E-05) | - | 0,987 |
| chr2:224257750:T:A (rs7578029) | *SERPINE2* | Putative causal | Infections of the skin and subcutaneous tissue | 7680/169219 | 1,12 (9,75E-05) | 0,72 (2,79E-02) | 0,967 |
| chr2:224497761:C:T (rs3738952) | *SERPINE2* | Putative causal | Malignant neoplasm of male genital organs | 4945/63465 | 0,86 (8,81E-05) | 0,93 (2,55E-01) | 0,580 |
| | | | Disorders of lens | 21994/154905 | 0,92 (5,70E-05) | 0,97 (4,16E-01) | 0,571 |
| chr5:150128981:C:G (rs2304058) | *PDGFRB* | Putative causal | Use of disulfiram, acamprosate or naltrexone | 1266/175633 | 1,18 (7,32E-05) | 1,05 (7,76E-01) | 0,943 |
| chr9:133271182:C:T (rs550057) | *ABO* | Putative causal | Intestinal infectious diseases | 16735/160164 | 0,95 (8,45E-05) | 1,00 (8,35E-01) | 0,567 |
| | | | Psoriatic arthropathies related co-morbidities | 37185/91355 | 0,95 (9,15E-05) | 1,00 (8,27E-01) | 0,555 |
| | | | Cholelithiasis, broad definition with cholecystitis | 15683/158425 | 0,94 (4,42E-06) | 0,98 (3,71E-01) | 0,465 |
| | | | Anaemias | 9145/50989 | 0,89 (4,69E-08) | - | 1,000 |
| | | | Other and unspecified anaemias | 4279/172180 | 0,91 (4,90E-05) | - | 1,000 |
| | | | Diseases of the blood and blood-forming organs and certain disorders involving the immune mechanism | 14375/162524 | 0,95 (2,86E-05) | 1,21 (4,33E-01) | 0,995 |
| | | | Other anaemias | 4337/172180 | 0,90 (2,61E-05) | - | 1,000 |
| | | | Diabetes medication | 25576/151276 | 0,95 (8,31E-05) | 0,99 (4,88E-01) | 0,513 |
| | | | Other (not insulin) diabetes medications | 21909/151323 | 0,95 (6,58E-05) | 0,98 (2,32E-01) | 0,513 |
| | | | Other diabetes, wide definition | 24662/144670 | 0,95 (9,45E-05) | 0,99 (4,72E-01) | 0,513 |
| | | | Endocrine, nutritional and metabolic diseases | 61193/115706 | 0,96 (5,56E-05) | 0,99 (2,81E-01) | 0,486 |
| | | | Pure hypercholesterolaemia | 6840/160461 | 0,91 (2,19E-06) | 0,98 (4,68E-01) | 0,555 |
| | | | Disorders of lipoprotein metabolism and other lipidaemias | 11042/160461 | 0,92 (1,28E-06) | 1,01 (7,41E-01) | 0,555 |
| | | | Metabolic disorders | 16438/160461 | 0,95 (2,36E-05) | 1,00 (9,57E-01) | 0,557 |
| | | | Cardiovascular diseases (excluding rheumatic etc) | 79685/97214 | 0,95 (5,57E-07) | 0,99 (3,32E-01) | 0,465 |
| | | | Diseases of veins, lymphatic vessels and lymph nodes, not elsewhere classified (FINNGEN) | 19930/97214 | 0,88 (2,29E-18) | 0,96 (4,96E-02) | 0,566 |
| | | | Other heart diseases | 43936/97214 | 0,95 (5,10E-07) | 0,98 (1,76E-01) | 0,483 |
| | | | Pulmonary heart disease | 3016/173883 | 0,87 (8,18E-07) | 1,01 (7,36E-01) | 0,513 |
| | | | Diseases of the eye and adnexa | 58498/118401 | 0,96 (9,42E-06) | - | 1,000 |
| | | | Visual field defects | 885/171027 | 0,81 (4,42E-05) | - | 1,000 |
| | | | Cardiovascular diseases | 86957/89942 | 0,96 (1,24E-06) | 0,98 (1,49E-01) | 0,465 |
| | | | Diseases of veins, lymphatic vessels and lymph nodes, not elsewhere classified | 22948/153951 | 0,90 (3,27E-17) | 0,97 (1,15E-01) | 0,566 |
| | | | DVT of lower extremities and pulmonary embolism | 6019/170880 | 0,82 (2,93E-20) | 1,00 (9,19E-01) | 0,553 |
| | | | Other heart diseases | 46991/129908 | 0,97 (8,84E-05) | 0,99 (6,08E-01) | 0,555 |
| | | | Phlebitis and thrombophlebitis (not including DVT) | 2506/153951 | 0,80 (6,27E-12) | 1,03 (5,24E-01) | 0,567 |
| | | | DVT of lower extremities | 3592/153951 | 0,77 (2,06E-21) | 0,99 (7,51E-01) | 0,553 |
| | | | Pulmonary embolism | 3016/173597 | 0,87 (8,65E-07) | 1,01 (7,28E-01) | 0,513 |
| | | | Pulmonary heart disease, diseases of pulmonary circulation | 3302/173597 | 0,88 (4,84E-06) | 1,03 (5,08E-01) | 0,514 |
| | | | Other embolism and thrombosis | 1416/153951 | 0,84 (2,26E-05) | 1,03 (5,96E-01) | 0,553 |
| | | | Varicose veins | 13928/153951 | 0,92 (1,30E-07) | 0,96 (8,50E-02) | 0,570 |
| | | | Other disorders of veins | 4588/153951 | 0,89 (1,63E-06) | 0,97 (4,31E-01) | 0,567 |
| | | | Venous thromboembolism | 6913/169986 | 0,83 (3,57E-22) | 1,03 (3,41E-01) | 0,567 |
| | | | Diseases of the blood and blood-forming organs and certain disorders involving the immune mechanism | 13899/163000 | 0,94 (1,20E-05) | 1,15 (5,60E-01) | 0,995 |
| | | | ILD-related co-morbidities | 85919/90980 | 0,97 (4,85E-05) | 0,99 (1,99E-01) | 0,513 |
| | | | ILD Co-morbidites, CVD and metabolic diseases | 62257/90980 | 0,96 (9,70E-06) | 0,98 (2,17E-01) | 0,555 |
| | | | Diseases of the circulatory system | 79046/97853 | 0,95 (2,06E-07) | 0,98 (1,94E-01) | 0,465 |
| | | | Chronic diseases of tonsils and adenoids | 19389/136923 | 0,94 (1,05E-05) | 0,99 (5,25E-01) | 0,553 |
| | | | Cholelithiasis | 15012/158425 | 0,93 (1,48E-06) | 0,98 (4,22E-01) | 0,566 |
| | | | Disorders of gallbladder, biliary tract and pancreas | 18474/158425 | 0,94 (5,21E-06) | 0,98 (3,20E-01) | 0,465 |
| | | | Endometriosis | 6502/57407 | 0,92 (1,00E-04) | 0,93 (1,62E-03) | 0,027 |
| | | | Co-morbidities of interest (NEURO) | 125910/50989 | 0,96 (2,18E-05) | 0,97 (1,73E-03) | 0,084 |
| | | | Other ILD-related CVD-co-morbidities | 3260/114642 | 0,86 (9,60E-08) | 1,01 (7,39E-01) | 0,514 |
| | | | Statin medication | 53518/123381 | 0,90 (4,19E-24) | 1,00 (8,35E-01) | 0,557 |
| | | | Diseases of the eye and adnexa | 53954/122945 | 0,97 (6,51E-05) | 1,00 (8,27E-01) | 0,995 |
| | | | Diseases of the ear and mastoid process | 31835/145064 | 0,96 (4,78E-05) | 0,98 (3,71E-01) | 0,995 |
| chr10:63288546:C:A (rs7080386) | *JMJD1C* | Putative causal | Cholelithiasis, broad definition with cholecystitis | 15683/158425 | 1,05 (7,02E-05) | 1,01 (6,71E-01) | 0,648 |
| | | | Cholelithiasis | 15012/158425 | 1,06 (4,07E-05) | 1,01 (6,75E-01) | 0,648 |
| chr15:101339772:G:A (rs111482836) | *PCSK6* | Putative causal | Anomalies of pupillary function | 169/175459 | 1,65 (6,99E-05) | 1,21 (2,26E-01) | 0,407 |
| chr19:55032292:G:A (rs199588110) | *GP6* | Putative causal | Benign neoplasm of meninges | 934/175965 | 6,40 (4,94E-05) | - | 1,000 |
| | | | Benign neoplasm of meninges, all cancers excluded | 934/147003 | 5,80 (7,73E-05) | - | 1,000 |

**Supplementary Table 8: Disease associations of the 19 putative causal variants and 13 functional variants in FinnGen.**
